# Functional analysis of the *teosinte branched 1* gene in the tetraploid switchgrass (*Panicum virgatum L.*) by CRISPR/Cas9-directed mutagenesis

**DOI:** 10.1101/2020.05.23.112961

**Authors:** Yang Liu, Weiling Wang, Bing Yang, Christopher Currey, Shui-zhang Fei

## Abstract

Tillering is an important biomass yield component trait in switchgrass (*Panicum virgatum L*.). *Teosinte branched 1* (*tb1*)/*Branched 1* (*BRC1*) gene is a known regulator for tillering/branching in several plant species; however, its role on tillering in switchgrass remains unknown. Here, we report physiological and molecular characterization of mutants created by CRISPR/Cas9. We successfully obtained non-chimeric *Pvtb1a* and *Pvtb1b* mutants from chimeric T0 mutants using nodal culture. The biallelic *Pvtb1a-Pvtb1b* mutant plants produced significantly more tillers and higher fresh weight biomass than the wild-type plants. The increased tiller production in the mutant plants resulted primarily from hastened outgrowth of lower axillary buds. Increased tillers were also observed in transgene-free T1 monoallelic mutants for either *Pvtb1a-Pvtb1b* or *Pvtb1b* gene alone, suggesting *Pvtb1* genes act in a dosage-dependent manner. Transcriptome analysis showed 831 genes were differentially expressed in the *Pvtb1*a-*Pvtb1b* double knockdown mutant. Gene Ontology analysis revealed downregulation of *Pvtb1* genes affected multiple biological processes, including transcription, flower development, cell differentiation, and stress/defense responses in edited plants. This study demonstrates that *Pvtb1* genes play a pivotal role in tiller production as a negative regulator in switchgrass and provides opportunities for further research aiming to elucidate the molecular pathway regulating tillering in switchgrass.

**Highlight:** Solid non-chimeric mutants were successfully isolated from CRISPR/Cas9-induced chimeric mutants using nodal culture. *Teosinte branched 1* (*tb1*) genes are involved in various pathways to regulate tillering in switchgrass.

## Introduction

Switchgrass (*Panicum virgatum*), a C4 perennial grass with demonstrated high biomass yield is native to North America and is well adapted to marginal land not suitable for food crops (Mitchell et al., 2008; Narasimhamoorthy et al., 2008). The low production cost and high lignocellulose-based biofuel potential make it well-suited for bioenergy crop (McLaughlin and Kszos, 2005). It was named the model species for herbaceous bioenergy crop by the U.S. Department of Energy in 1991 following more than a decade of research (Wright and Turhollow, 2010). Switchgrass is an out-crossing species with varied ploidy levels and is self-incompatible with individual plants being highly heterozygous (Narasimhamoorthy et al., 2008). Upland and lowland ecotypes are the two major representative taxa of switchgrass (Zhang et al., 2011). Most of the low-tillering lowland ecotypes are tetraploid (2n=4x=36), while the high-tillering upland ecotypes contain both tetraploids and octoploids (2n=8x=72) with hexaploids (2n=6x=54) being reported rarely (Zhang et al., 2011). Unlike the lowland ecotype, which has a caespitose growth habit and poor freezing tolerance, upland ecotypes are highly rhizomatous and cold hardy (Mitchell and Schmer, 2012).

High biomass yield is a high priority for switchgrass breeding (Casler, 2010; Okada et al., 2010; Lu et al., 2013). Tiller number is positively correlated with biomass yield (Das et al., 2003; Boe, 2007; Boe and Beck, 2008). Das et al. (2003) reported tiller number contributes directly to biomass yield of switchgrass. Therefore, understanding the molecular mechanism regulating tillering in switchgrass can facilitate development of high-yielding cultivars. Tillers result from the outgrowth of axillary buds at nodes in grass species and the number of tillers produced from a single plant varies greatly by species and genotypes. Tillering or branching is regulated by numerous endogenous and environmental factors in both eudicots and monocots (Kebrom et al., 2010; Whipple et al., 2011; Reddy and Finlayson, 2014; Gonzalez-Grandio et al., 2017; Holalu and Finlayson, 2017). For instance, auxin is a well-known contributor to apical dominance, which inhibits axillary bud outgrowth, while cytokinins (CKs) promote bud outgrowth (Thimann and Skoog, 1933; Morris, 1977; Minakuchi et al., 2010; Braun et al., 2012). Strigolactones (SLs), another key plant hormone, represses branching or tillering in both monocots and eudicots (Sorefan et al., 2003; Zou et al., 2006; Arite et al., 2007). Additionally, tiller production is often inhibited under severe shade (Kebrom et al., 2006; Finlayson et al., 2010; Kebrom et al., 2010), insufficient mineral nutrients or photosynthates (Wang et al., 2019).

The *teosinte branched 1* (*tb1*) gene is an important tillering/branching-related transcription factor gene that regulates tillering by integrating environmental and developmental cues (Doebley et al., 1997; Whipple et al., 2011; Seale et al., 2017). In cultivated maize (*Zea mays* L.), an insertion in the regulatory region of the *tb1* gene elevates its expression at leaf axils where axillary buds are located and greatly suppresses tillering, while its ancestor teosinte lacking the insertional sequence is highly branched (Studer et al., 2011). *tb1* belongs to the *TCP* (*TEOSINTE BRANCHED1*, *CYCLOIDEA*, *PCF*) gene family that all encode proteins with a 59-amino acid, non-canonical basic helix-loop-helix (bHLH) motif that allows DNA binding and protein-protein interactions (Martin-Trillo and Cubas, 2010). Closely related *tb1* genes with similar functions have been identified in both monocots and dicots (Kebrom et al., 2006; Aguilar-Martinez et al., 2007; Martin-Trillo et al., 2011; Choi et al., 2012; Nicolas et al., 2015). For example, the *OsTB1* gene functions as a negative regulator for lateral branching in rice (*Oryza sativa* L.) (Takeda et al., 2003; Choi et al., 2012). The ortholog of *tb1* in bread wheat (*Triticum aestivum* L.) regulates inflorescence architecture and outgrowth of axillary buds (Dixon et al., 2018). In Arabidopsis, two orthologs of *tb1*, *BRANCHED 1* (*BRC1*) and *BRANCHED 2* (*BRC2*), regulate branching with *BRC1* having the major effect (Aguilar-Martinez et al., 2007). Many studies demonstrate *tb1* genes integrate multiple signaling pathways to regulate bud outgrowth and tillering (Kebrom et al., 2013; Rameau et al., 2015). Low ratio of red light to far-red light (R/FR), as a result of shade, inhibits the outgrowth of axillary buds in sorghum (*Sorghum bicolor* (L.) Moench) by upregulating the expression of *SbTB1*, an ortholog of *tb1* (Kebrom et al., 2006; Kebrom et al., 2010). In Arabidopsis (*Arabidopsis thaliana* (L.) heynh.), *BRC1* suppresses lateral bud growth in response to shade by promoting abscisic acid accumulation inside axillary buds (Gonzalez-Grandio et al., 2017). Cytokinins activate axillary bud outgrowth by inhibiting localized *PsBRC1* expression in pea (*Pisum sativum* L.) (Braun et al., 2012) or *OsTB1* in rice (Minakuchi et al., 2010). These studies all demonstrate that *BRC1/TB1* is a common target for hormonal and environmental signals that regulate tillering/branching across species. Hence, understanding the function of switchgrass *tb1* genes can provide valuable information to the understanding of the tillering mechanisms in switchgrass.

The lowland switchgrass cv. Alamo from which the reference genome was produced is an allotetraploid (2n = 4x = 36, NNKK). It is therefore hypothesized that the majority of genes are present as homoeologs. There are two *Pvtb1* genes (*Pvtb1a* and *Pvtb1b*) in ‘Alamo’ with 89%. DNA sequence identity between them (*Panicum virgatum* v4.1, DOE-JGI, http://phytozome.jgi.doe.gov/). Because of its self-incompatibility, it is difficult to obtain homozygous mutants in switchgrass by inbreeding hemizygous mutants as is done for self-pollinating transgenic plants. The clustered regularly interspaced short palindromic repeat (CRISPR)/CRISPR-associated protein 9 nuclease (Cas9) based genome editing tools have become a powerful genetic tool for gene function analysis or crop improvement (Zhao et al., 2016; Wang et al., 2018; Yang et al., 2018). A multiplex CRISPR/Cas9 platform can be used to simultaneously edit multiple genes, which is particularly advantageous for a polyploid species such as switchgrass in which homeologs of a same gene exist, or for members of a gene family that are present in tandem (Zhao et al., 2016).

We previously created switchgrass *tb1* mutant plants by using CRISPR/Cas9 (Liu et al., 2018). However, the allelic composition of *Pvtb1* genes in these T0 mutant plants was not fully characterized. In addition, it remained possible that these primary mutants are chimeric, which prevented us from accurately assessing the function of *tb1* genes. In plant species that can be vegetatively propagated, non-chimeric, solid mutants can be successfully isolated from chimeric mutants (Maluszynski et al., 1995). For example, separating mutated sectors from chimeric mutants by *in vitro* multiplication has been achieved in banana, cassava and other vegetatively propagated crops (Harten et al., 1981; Novák et al., 1990; Maluszynski et al., 1995; Das et al., 2000). Switchgrass can be readily propagated using node culture (Alexandrova et al., 1996). Hence, generation of primary mutants with CRISPR/Cas9, followed by nodal culture would allow us to generate non-chimeric mutants without the need of producing progeny.

The relative ease with which transgene-free mutant plants can be obtained is a unique advantage for CRISPR/Cas9-based genome editing. Transgene-free mutants have been obtained in multiple crops using CRISPR/Cas9-based genome editing tools and stable inheritance of CRISPR/Cas9 induced mutations has been demonstrated in other plant species, (Pyott et al., 2016; Zhang et al., 2016; He et al., 2018) but not yet in switchgrass. Germplasm generated through CRISPR/Cas9-based genome editing method without foreign DNAs may bypass or simplify the regulatory process required for crops created with traditional transgenic approaches (Waltz, 2018).

Here, we report the successful use of micropropagation to generate non-chimeric mutants from chimeric primary mutants induced by CRISPR/Cas9, eliminating the need of obtaining progeny mutants for gene function analysis. Moreover, the transmission of CRISPR/Cas9-induced mutations in switchgrass is demonstrated in this study. Transgene-free progeny mutants, preserving the phenotypic effect of *Pvtb1* mutations similar to that exhibited in T0 mutants, were generated in this study. We showed that double biallelic mutant for *Pvtb1a* and *1b* genes enhanced tiller production and increased biomass yield, indicating that *Pvtb1* genes negatively regulate tillering in switchgrass. Transcriptome analysis of a *Pvtb1* knockdown mutant 52-1 and wild-type (WT) plant WT-1 suggest that *Pvtb1* genes are involved in multiple pathways to regulate tillering in switchgrass.

## Materials and methods

### Micropropagation and seed propagation of CRISPR/Cas9-induced *Pvtb1* mutants

Primary mutant plants were generated previously (Liu et al., 2018). Briefly, the CRISPR/Cas9 construct containing two gRNAs targeting the conserved regions of *Pvtb1a* and *Pvtb1b* genes was delivered into the genome of switchgrass through *Agrobacterium*-mediated transformation of caryopsis-derived embryogenic callus from which primary mutant plants were obtained. To produce non-chimeric mutants from chimeric primary mutants, we micropropagated *Pvtb1* mutants by culturing nodes containing axillary buds according to (Harten et al., 1981; Alexandrova et al., 1996). Each of the two basal nodes from a stem containing an axillary bud with a portion of (~1.5cm long) of internode above and below the node was selected for *in vitro* culture on the MS-0 medium with 3 gL^−1^ Phytagel (Sigma Chemical Co., St. Louis) and no plant growth regulator. These nodal segments were surface sterilized in commercial bleach (5% aqueous solution of sodium hypochlorite) for 30 min, and then rinsed three times with sterile distilled deionized water. Three experiments were conducted for the 52-1 and 35-2 mutants and their corresponding WT (WT-1) plants with 20 nodes for each experiment. After 8 weeks of culture in a growth chamber with a light intensity of 140 μmol m^−2^ s^−1^ at a photoperiod of 16/8 hours light/dark and a temperature of 25 °C, regeneration efficiency (regenerated plantlets/total nodes number) were calculated. After roots emerged, plantlets were grown in 6-inch diameter (1L) pots using commercial soil mix (Sunshine soil mix #1, Sun Gro) of peat moss and perlite and moved into a mist room for 7-10 days before they were placed in a greenhouse at 26 °C with a 16/8h (day/night) photoperiod and a light intensity of approximately 400 μmol m^−2^ s^−1^.

To determine the inheritance of CRISPR/Cas9 induced mutations in switchgrass, T1 progeny were obtained by crossing mutants with genetically compatible WT plants. Seeds were harvested from the mutant parents and were sowed in a 1020 tray (54.5 cm L × 27.8 cm W × 6.2 cm H) with the same soil as described previously that was maintained at a mist room until germination. Seedlings were transferred to the same greenhouse as described earlier. The presence of the transgene was analyzed using PCR with gRNA/Cas9-specific primers (Table S6).

### Genotyping of CRISPR/Cas9-induced *Pvtb1* mutants by Next Generation Sequencing

NGS was used to genotype the *Pvtb1* mutant plants generated from micropropagation of primary mutants and T1 *Pvtb1* mutant progeny. To amplify the target fragments of each *Pvtb1* gene, gene-specific primers were designed for *Pvtb1a* and *Pvtb1b* respectively. The amplicon size for *Pvtb1a* is 250 bp, while that of *Pvtb1b* is 288 bp, both are sufficiently long to cover the two target sequences. The Illumina overhang adapter sequences were added to the gene-specific primers (Table S6). A second round of PCR was used to add the dual indices with the Nextra XT Index Kit (Illumina, San Diego, CA, USA). The quality of these sequencing libraries was determined by the Qubit® 2.0 Fluorometer. The 150-cycle HiSeq sequencing was done for all the amplicon libraries at the DNA Facility at Iowa State University (ISU). For each library, at least 5,000 reads were generated to determine the sequence of the respective amplicons. The results were analyzed by CRISPR-DAV pipeline (Wang et al., 2017).

### Sequence alignment and phylogenetic analysis

The sequence of maize *tb1* gene was used to blast the sequences of *Pvtb1* genes from the *P. virgatum* genome project v4.1 (Panicum virgatum v1.0, DOE-JGI, http://phytozome.jgi.doe.gov/). The *Pvtb1* genes were subsequently isolated and sequenced with the *Pvtb1* gene-specific primers (Table S6). Full-length putative amino acid sequences of *tb1* gene orthologs were obtained from Phytozome (https://phytozome.jgi.doe.gov/pz/portal.html) (Table S7). Alignments of the putative protein sequences of TB1 were performed using Clustal Omega (Sievers et al., 2011) (https://www.ebi.ac.uk/Tools/msa/clustalo/).

### Phenotypic characterization of *Pvtb1* mutant plants with hydroponic culture

To investigate the effects of *Pvtb1* genes on phenotype of switchgrass, we established a hydroponic system to observe plant growth. Nodal segments were cut from the wild-type plant (WT, genotype AABB) and mutant 52-1-3 (genotype aabb), 52-1-1 (genotype Aabb) plants, and cultured *in vitro* as described previously. Plants of similar height that were grown in 6-inch pots for 10 days were transferred into a hydroponic device in a greenhouse where temperature was maintained at 26°C with a 16h/8h (day/night) photoperiod with a light intensity of approximately 400 μmol m^−2^ s^−1^. For hydroponic culture, the stem base of each plant was wrapped around by sponge and placed into a small basket which was then inserted into a hole on the lid of a plastic container (51 cm L × 43 cm W × 15 cm H) that was filled with a nutrient solution (16 - 4 - 17 Oasis Hydroponic, JR Peters Inc., PA, USA). Electrical conductivity (EC) and pH of the nutrient solution were measured by a portable pH/EC/Temperature meter (HI9813-6, Hanna Instruments, Inc., RI, USA) daily and maintained at 1.35 mS cm^−1^ and 6.0, respectively. The pH was lowered or raised by phosphoric acid or sodium hydroxide, respectively, while the EC was maintained by adding concentrated fertilizer stock solution or clear water. The nutrient solution was aerated by an aquarium air pump (ActiveAQUA, Hydrofarm, CA, USA) to ensure adequate oxygen supply.

### Statistical analysis of morphological traits

Primary, secondary, tertiary or quaternary tillers were visually determined according to Davidson & Chevalier (1987). Primary tillers were defined as those which arise from leaf axils in nodes of the main stem whereas secondary tillers arise from leaf axils in nodes of the primary tillers. By the same rule, tertiary and quaternary tillers were defined as those arise from leaf axils of nodes of secondary and tertiary tillers, respectively. Adventitious roots of newly formed tillers tend to have creamy white color whereas roots of older tillers appear in darker color. This color variation was used to assist classification of tillers. Plant height was determined by measuring the tallest tiller for each plant. Root length was determined by measuring the length of the longest root for each plant. Stem diameter was determined by the average diameter of the middle internode of five largest tillers for each plant. The number of tillers and roots (longer than 2 cm) were also counted. To determine biomass yield, pot-grown plants were manually cut at the ground level using a pruning shears and weighted for fresh weight. Samples were then oven-dried at 70°C for 4 days when they reached a constant weight to obtain the dry weight. Three to six biological replicates were used in each measurement. Student’s t-test was used to determine the significance of difference between the mutant plants and the WT plants.

### Transcriptome sequencing and analysis of the mRNA-seq data

It was shown in the expression database that *Pvtb1* genes in switchgrass expressed highly in axillary buds (Panicum virgatum v1.0, DOE-JGI, http://phytozome.jgi.doe.gov/). Thus, we extracted RNAs from axillary buds of tillers. The mutant 52-1 with highly enhanced tiller production and the WT plant (WT-1) generated from the same callus line from which the mutant 52-1 was obtained were chosen for mRNA-seq. Three biological replications were sampled for each plant. All visible axillary buds collected from two to three tillers are treated as one biological replicate. Total RNA was isolated from tissue samples using the Qiagen RNeasy Plant Isolation kit according to the manufacturer’s protocol (Qiagen Inc., Valencia, CA, USA) and the RNA quality was checked using BioAnalyzer 2100 (Agilent Technologies, Santa Clara, CA, USA). cDNA libraries were constructed by the ISU DNA Facility according to the instructions in the Illumina sequencing manual. The Illumina HiSeq 3000 150 paired-end platform was used for sequencing by the DNA Facility at Iowa State University (http://www.dna.iastate.edu/).

Library adapter and low quality nucleotides were trimmed off using Trimmomatic (Bolger et al., 2014). Then, STAR was applied to align the trimmed reads to the reference genome V4.1 of switchgrass (Dobin et al., 2013). The number of reads mapped to each gene was calculated using HTseq-count (Anders et al., 2015). Normalization was done using the Upper Quartile method and gene expression differences were analyzed with package edgeR (Robinson et al., 2010) in the statistical software ‘R’ (Version 3.4.3) (The R FAQ, https://CRAN.R-project.org/doc/FAQ/). Genes are classified as differentially expressed when an absolute log2-fold change value is ≥1 and a false discovery rate is ≤ 0.05 (Benjamini and Hochberg, 1995). Gene Ontology (GO) analysis for DEGs were performed using the Database for Annotation, Visualization and Integrated Discovery (DAVID) (Huang et al., 2009a; Huang et al., 2009b). ReviGO (Supek et al., 2011) and Cytoscape (Shannon et al., 2003) were applied to visualize enriched GOs.

## Results

### Micropropagation can effectively generate non-chimeric mutants from chimeric primary (T0) CRISPR/Cas9 - induced mutants

We micropropagated the primary (T0) *Pvtb1* mutants (52-1 and 35-2) using *in vitro* node culture (Figure S1). After about 8 weeks of culture, 48 and 20 plants were successfully regenerated from the primary mutants 35-2 and 52-1, respectively.

To determine the allelic compositions of *Pvtb1a* and *Pvtb1b* genes in 10 randomly selected micropropagated plants including 9 plants generated from primary mutants and 1 plant from the wild-type, sequencing libraries for the *tb1a* amplicons with an insert size of 250 bp and *tb1b* amplicons with an insert size of 288 bp were constructed for each individual plant. For each library, at least 5,000 reads were generated from Illumina HiSeq3000 to determine the sequence of the respective amplicons. Three distinct genotypes were found among the five micropropagated plants (52-1-1, -2, -3, -4, -5) derived from 52-1, while four distinct genotypes were observed among the four micropropagated plants (35-2-1, -2, -3, -4) derived from 35-2, clearly indicating that the two primary *Pvtb1* mutants from which micropropagated plants were obtained are chimeric (Table 1).

**Table 1.**
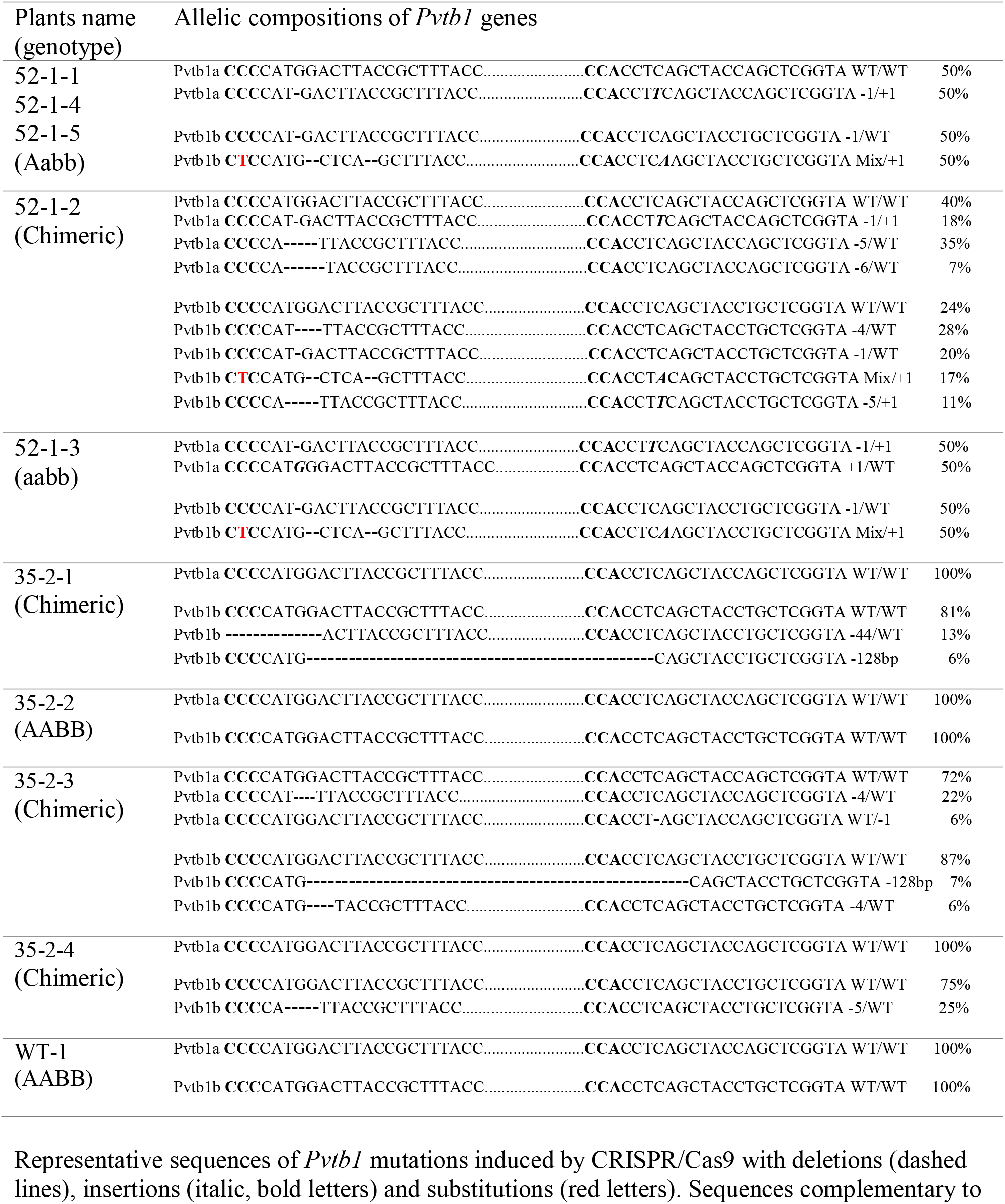
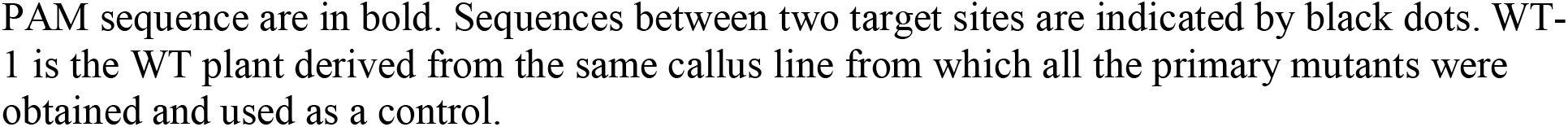
Estimation of allelic composition of *Pvtb1* genes in plants regenerated from node culture of the primary mutants 52-1 and 35-2.

Four non-chimeric, solid mutants, 52-1-1, 52-1-3, 52-1-4, and 52-1-5 were obtained from the T0 mutant 52-1 with 52-1-1, -4, and -5 all having the same genotype, i.e. 50% of WT *Pvtb1a* allele and 50% *Pvtb1a* allele with a G deletion at the first target site and a T insertion at the second target site (Table 1), suggesting these plants were likely originated from a single progenitor mutant cell carrying a monoallelic *Pvtb1a* mutation. In the other solid mutant 52-1-3, all alleles of the *Pvtb1a* were mutated with half of the *Pvtb1a* alleles contains a G deletion at the first target site and a T insertion at the second target site, while the remaining half of *Pvtb1a* alleles contains only one change, an insertion of G at the first target site (Table 1), suggesting 52-1-3 was likely derived from a single progenitor mutant cell carrying biallelic mutations of *Pvtb1a* gene. These four plants all contain the same biallelic mutations for *Pvtb1b* with no WT alleles (Table 1). Fifty percent of the amplicons from each of the four solid mutants contained one C deletion at the first target site, while the remaining half amplicons contained the same mixed mutation (deletions, insertions and substitution) at the first target site and one A insertion at the second target site (Table 1), suggesting these four plants were developed from a single progenitor cell or mutant sector carrying biallelic mutations of the *Pvtb1b* gene.

Some of the micropropagated plants remained chimeric following micropropagation. In 52-1-2, four types of *Pvtb1a* amplicons including 3 different mutant alleles accounting for 60% of the reads and 40% WT reads were found (Table 1). In 35-2-3, there were three types of *Pvtb1a* amplicons including 28% mutated amplicons and 72% WT (Table 1). For *Pvtb1b*, in 52-1-2, 35-2-1, 35-2-3, and 35-2-4, the percentage of mutant reads are 76%, 19%, 13%, and 25%, respectively (Table 1). The makeup of the WT and mutant alleles is not proportional to the expected allelic composition for a tetraploid species, strongly indicating these four micropropagated plants (52-1-2, and 35-2-1, -3, -4) remain chimeric mutants comprising a mixture of mutant cells carrying different mutations induced by CRISPR/Cas9. These results indicate the axillary bud contained in the explants used for node culture were likely chimeric.

### Knock-out of *Pvtb1* genes leads to increased tillering in switchgrass

Based on the most recent annotated switchgrass genome in Phytozome (Goodstein et al., 2012) (https://phytozome.jgi.doe.gov/pz/portal.html), two *tb1-like* genes, *Pvtb1a* (Pavir.Ia00838/Pavir.9NG142700) and *Pvtb1b* (Pavir.Ib04362/Pavir.9KG031700), were isolated from switchgrass using gene-specific primers. These two *Pvtb1* genes reside on homeologous chromosomes 9N and 9K, thus are likely homeologous to each other. Amino acid sequence alignment of *Pvtb1* gene products and other *tb1* orthologs of related species showed that both PvTB1 proteins have the conserved TCP domain and R domain (Figure 1). To determine the function of *Pvtb1* genes, tiller numbers of the *Pvtb1a-Pvtb1b* double biallelic mutants (*aabb*), *Pvtb1a* monoallelic and *Pvtb1b* biallelic mutant plants (*Aabb*) and WT (AABB) plants were compared. Because switchgrass tillers develop from nodes near or below soil surface, we grew plants in a hydroponic system, which allows close monitoring of tiller and root growth without the destructive excavation of the underground parts of these plants. As shown in Figure 2, the first primary tiller of WT plants did not develop until the end of the fourth week following establishment in the hydroponic system, while most of the mutant plants developed their first primary tiller by the end of the first week. After 4 weeks, the average tiller number of *aabb* mutants is 5.7 which is about 2 times the tiller number of WT plants, while the average tiller number of *Aabb* mutants is 3.7 which is about 1.5 times the tiller number of WT plants. The average tiller number of *aabb* mutants is not significantly higher than the average tiller number of *Aabb* mutants, suggesting the *Pvtb1a* gene has a minor effect on tillering in switchgrass.

**Figure 1.**
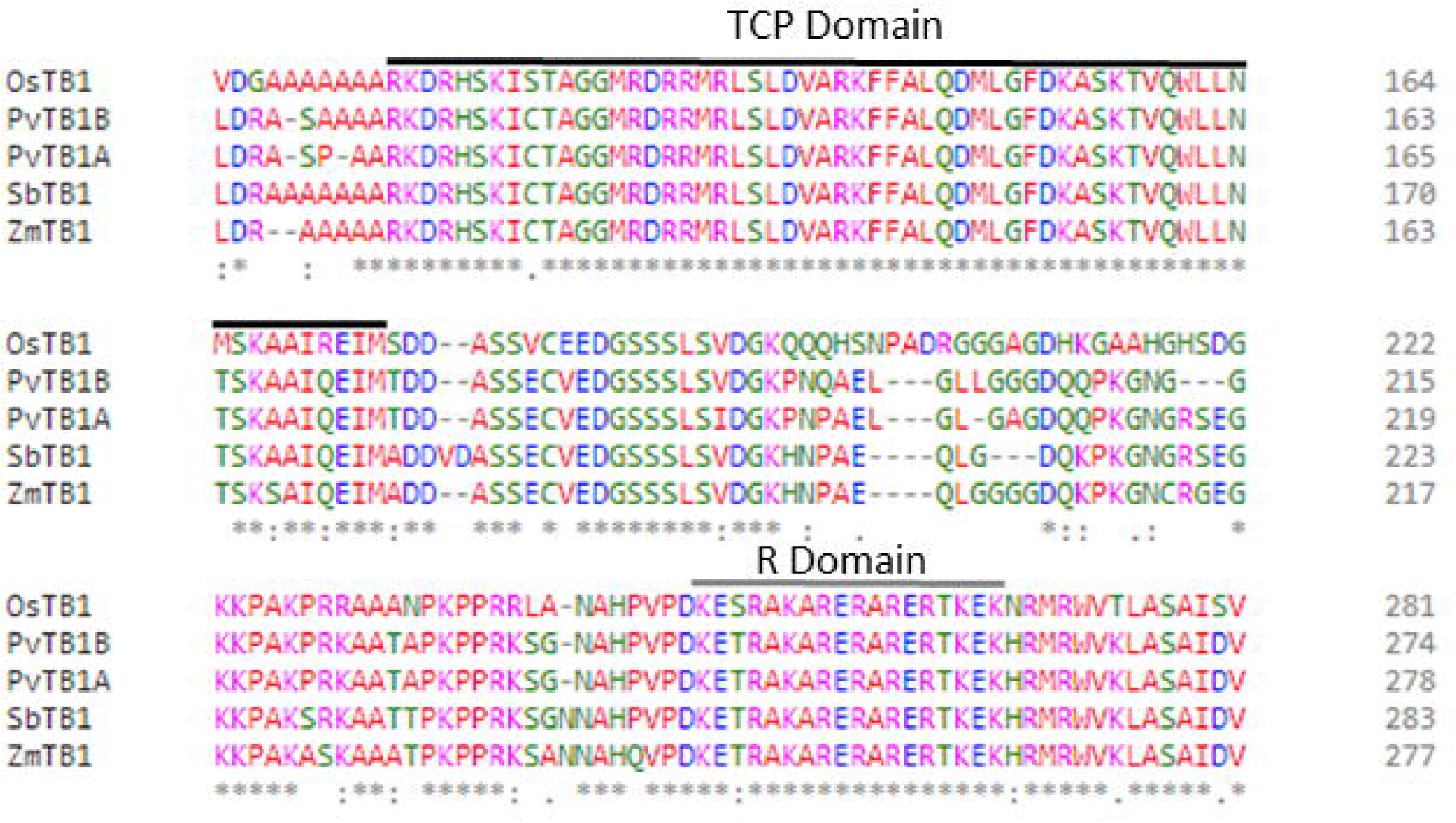
Multiple sequence alignment of TB1 proteins. (A). Multiple sequence alignments of TB1 orthologs from various species. The black line indicates the TCP domain, while the grey line indicates the R domain. Protein names are shown before each sequence.

**Figure 2.**
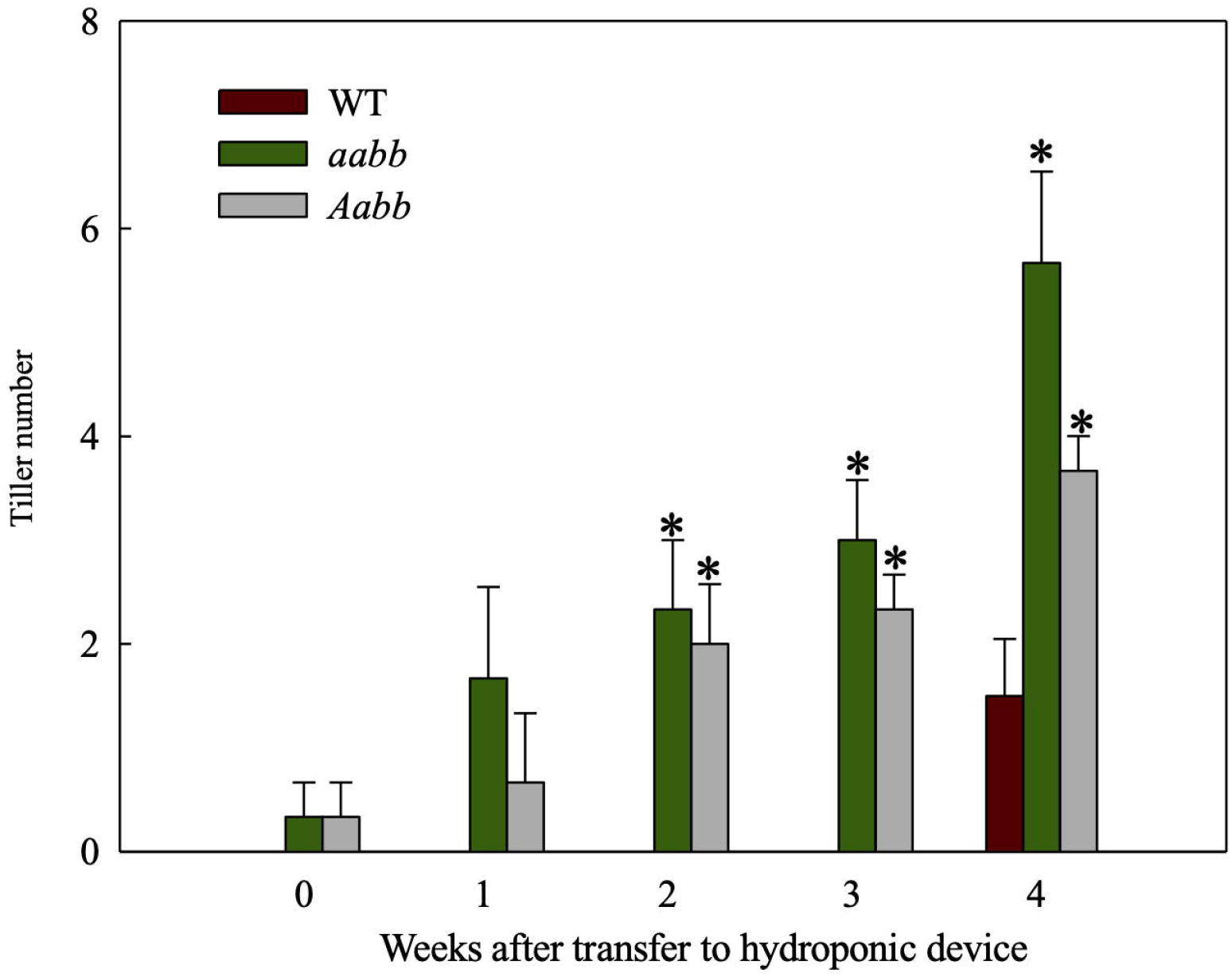
Tiller development in the double biallelic mutant 52-1-3 (*aabb*) and 52-1-1 mutant (monoallelic for *Pvtb1a* and biallelic for *Pvtb1b, Aabb*) and the WT at various times after transfer to hydroponic devices. Values are means ± s.d. (mutants, n = 3 plants; WT, n = 6 plants). * indicated significance differences between mutants and WT at P < 0.05.

To further investigate where the increased tiller originate from mutants, development of tillers of different orders in the *aabb* mutant plants and WT plants were closely monitored for 8 weeks in a separate hydroponic experiment. As shown in Figure 3A, the *pvtb1a-pvtb1b* (aabb) mutant plants had significantly more tillers than WT plants. The change of tiller numbers over the course of 8 weeks show the tiller numbers of mutant plants are significantly higher than WT plants at all weekly sampling times (Figure 3B). The WT plants did not generate new tillers until the end of the fourth week after establishment in the hydroponic system, while the mutant plants started to produce new tillers at the end of the first week, which is similar to the results from the shorter-term study described above. However, the *pvtb1a-pvtb1b* mutant plants had no significant effect on the number of primary tillers (1°) compared with WT plants (Figure 3C and D), which suggested that *Pvtb1* genes do not regulate the formation of primary axillary buds in the main stem, and the significant increase in secondary (2°), tertiary (3°), or quaternary (4°) tillers in the mutant plants is the result of hastened outgrowth of primary axillary buds. To further confirm the effect of *Pvtb1* on the outgrowth of axillary buds and eliminate possible effect of physiological conditions of the starting materials, primary tillers developed from the *pvtb1a-pvtb1b* double biallelic mutant plants and the WT plants were grown as starting material in a separate hydroponic study. The total number of new tillers developed from the primary tillers of the mutant plants was three-fold of that produced by the primary tillers of the WT plants (Figure 4A and B). Similarly, plants established from the primary tillers of the *pvtb1a-pvtb1b* mutant had similar number of the first order tillers (1°) to that of the WT, but had greatly increased number of secondary (2°) and tertiary tillers (3°) than those from the primary tillers of the WT. These results strongly suggest that, regardless of the origin of the starting materials, the *pvtb1a-pvtb1b* mutant plants consistently promote axillary bud outgrowth, increasing tiller number. Aerial tillers, however, were not observed in either the mutant plants or the WT plants, suggesting that *Pvtb1* genes did not affect the outgrowth of the upper axillary buds. Interestingly, while the first axillary bud (the lowest in the stem base) in the WT was generally dormant, it was frequently elongated to eventually become a tiller in the *pvtb1a-pvtb1b* mutant (Figure S2). These results further demonstrate that *Pvtb1* genes affect the outgrowth of the axillary bud rather than the formation of new axillary buds.

**Figure 3.**
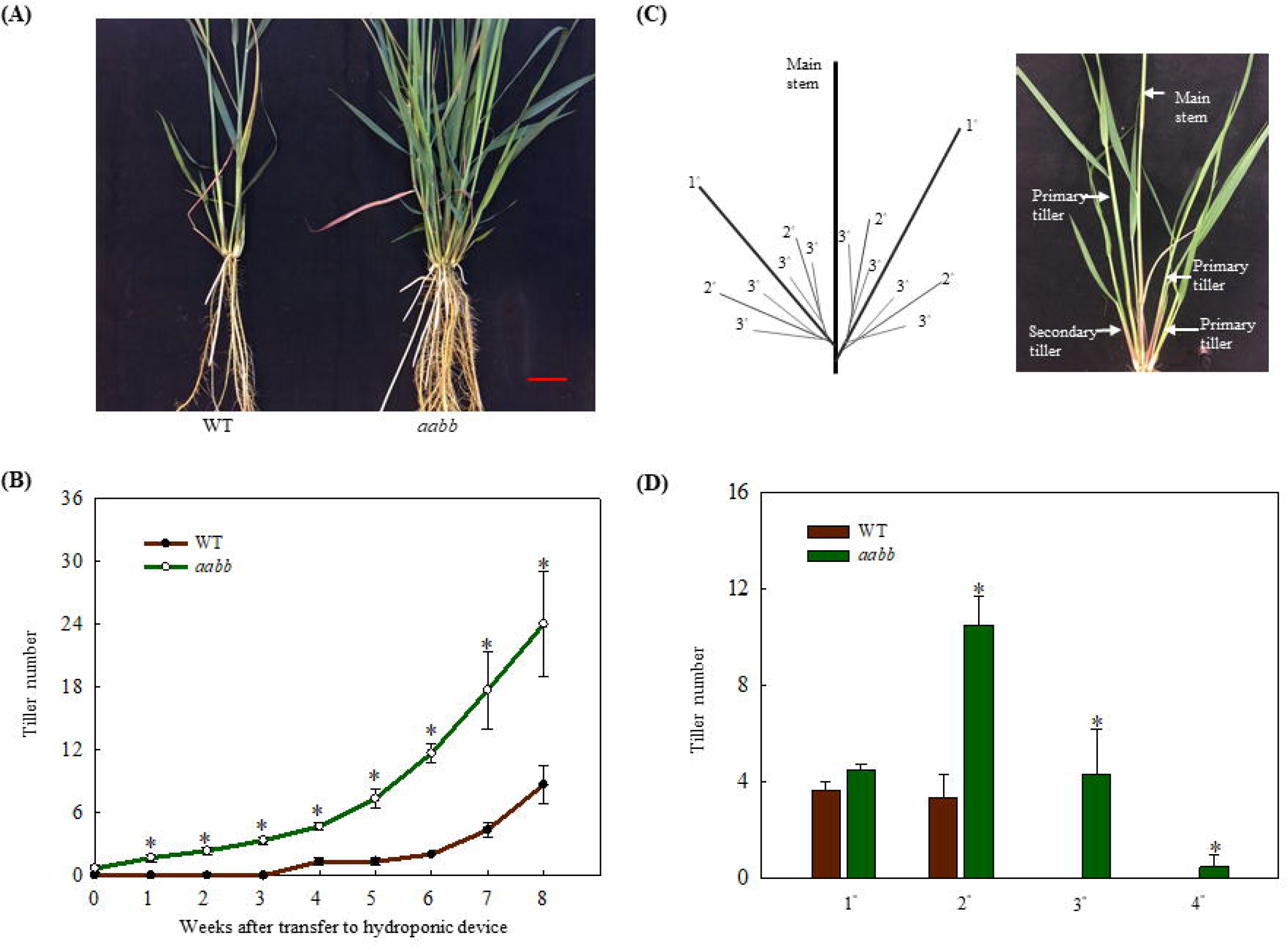
Phenotypic characterization of the *Pvtb1a-Pvtb1b* biallelic mutant (52-1-3, *aabb*) and the WT. (A). Phenotype of the *Pvtb1a-Pvtb1b* biallelic mutant and the WT after 8 weeks of growth in a hydroponic device. Bar = 3 cm; (B). Weekly changes in tiller numbers in the 52-1-3 mutant and the WT after transfer to a hydroponic device; (C). Schematic diagram for tiller ordering in switchgrass. 1° denotes primary tillers, 2° denotes secondary tillers, 3° denotes tertiary tillers; (D). Number of tillers of different orders in the 52-1-3 mutant and the WT after 8 weeks of growth in hydroponic devices. Values are means ± s.d. (B, n = 3 plants; D, n = 6 plants). * indicated significant differences at *P* < 0.05.

**Figure 4.**
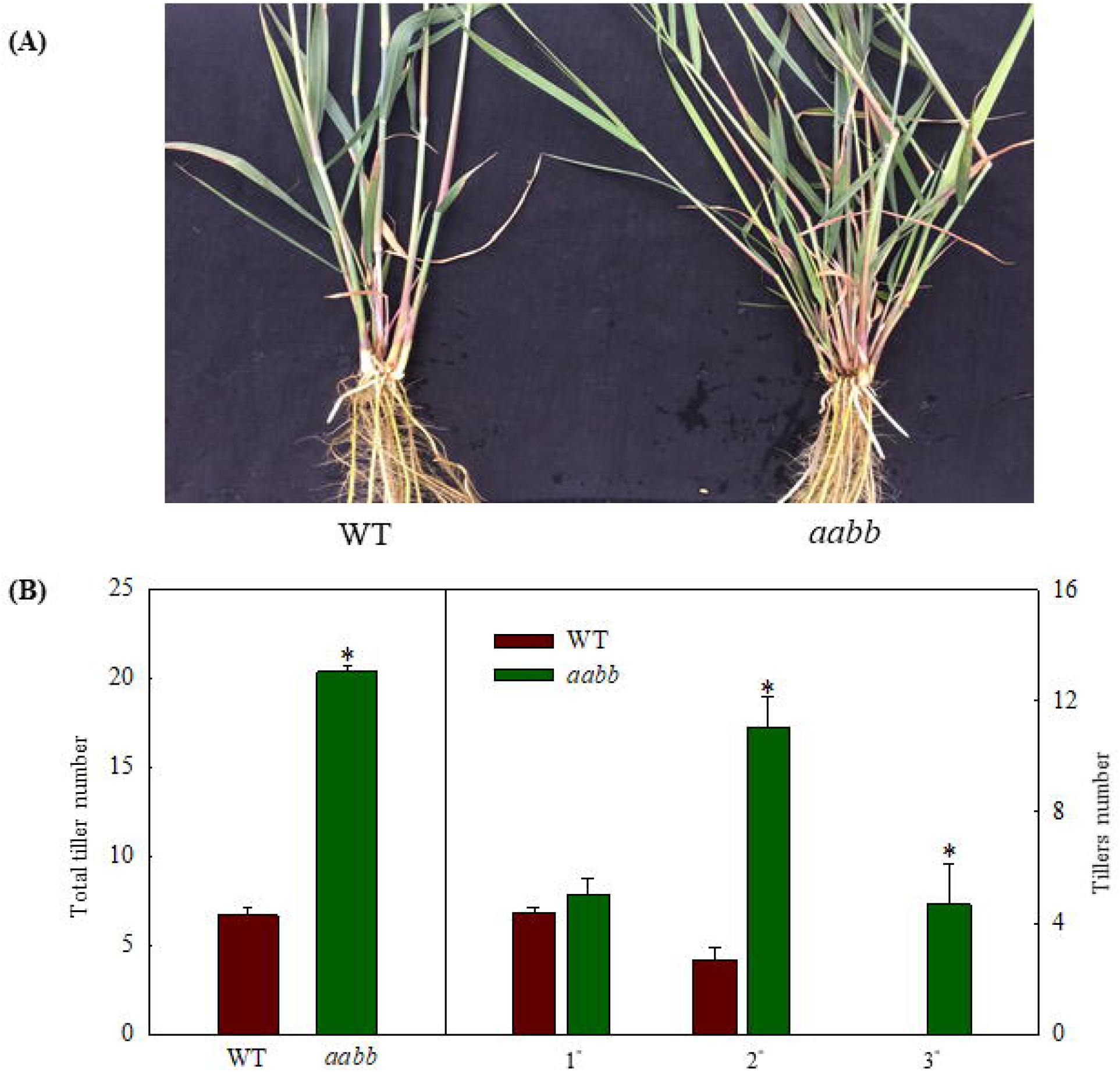
Phenotypic characterization of tiller production with primary tillers developed from main stems of the *Pvtb1a-Pvtb1b* double biallelic mutant (52-1-3, *aabb*) as starting materials and cultured in a hydroponic devise for 8 weeks. (A). Visual appearance of the WT (left) and the mutant (right) after 8 weeks of culture; (B). Total number of tillers and tiller number for each class of tillers after 8 weeks of culture. 1° denotes primary tillers, 2° denotes secondary tillers, 3° denotes tertiary tillers. Values are means ± s.d. (n = 3 plants). * indicated significant differences at *P* < 0.05.

### *Pvtb1* knock-out mutant plants exhibited increased root production and biomass yield

The average root number in mutant plants was similar to that of the WT plants at the beginning of the hydroponic culture (Figure 5). Starting from the second week, the difference between the average root number in mutant plants and WT plants started to increase as the experiment progressed (Figure 5). At the end of the experiment, the average root number of the mutant plants was 28, twice the average root number of the WT plants (Figure 5). Hence, disrupting *Pvtb1* gene function in switchgrass enhances root production.

**Figure5.**
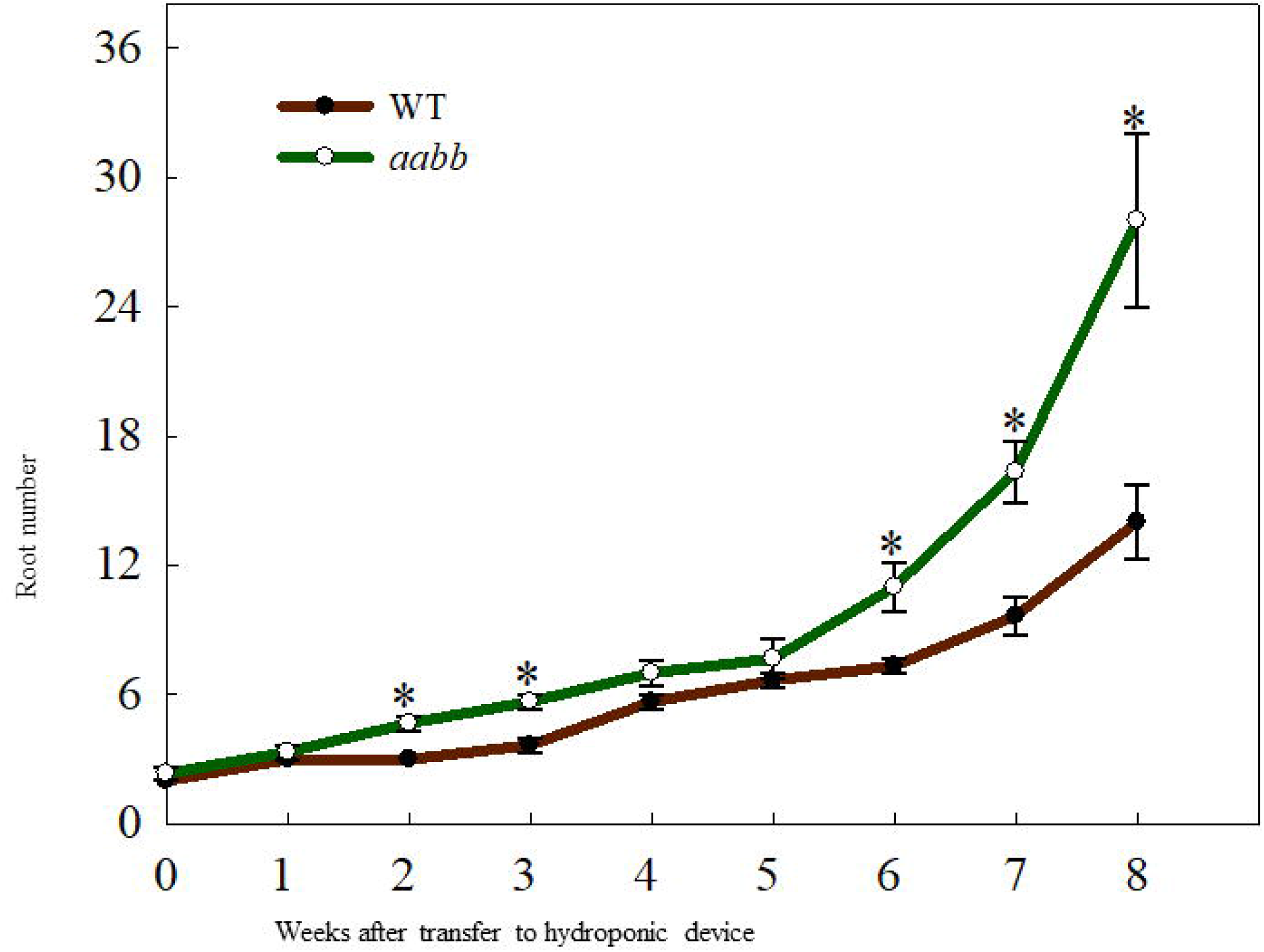
Weekly changes in root number in the *Pvtb1a-Pvtb1b* double biallelic mutant (52-1-3, *aabb*) and the WT after transfer to hydroponic devices. Values are means ± s.d. (B and D, n = 3 plants; C, n = 6 plants). * indicated significant differences at *P* < 0.05.

To eliminate the potential unintended effect of the hydroponic growth environment on growth and development, we also evaluated tiller number, stem diameter, plant height, fresh weight and dry weight of *pvtb1a-pvtb1b* mutant plants and WT plants grown in the soil. Similar to the hydroponic experiments, *pvtb1a-pvtb1b* mutant plants produced significantly more tillers (2.53-fold) compared with the WT plants (Table 2). Plant height of the mutant plants was similar to that of the WT, while stem diameter of the mutant plants was 13.6% smaller than that of the WT (Table 2). The fresh and dry biomass of the mutant plants increased 29.6% and 15.5% over these of the WT plants, respectively, suggesting the enhanced numbers of tillers of mutants were more than sufficient to compensate for the loss in biomass from decreased stem diameter (Table 2).

**Table 2.** Phenotype of the WT and the micropropagated biallelic mutant plants, after 12 weeks of growth in pots filled with soil.

| Genotype | Tiller number | Stem diameter (mm) | Plant height (m) | Fresh weight (g plant<sup>-1</sup>) | Dry weight (g plant<sup>-1</sup>) |
| --- | --- | --- | --- | --- | --- |
| AABB (WT) | 22.4 ± 1.44<sup>b</sup> | 4.56 ± 0.13<sup>a</sup> | 1.67 ± 0.11<sup>a</sup> | 34.5 ± 1.30<sup>b</sup> | 10.3 ± 0.98<sup>a</sup> |
| *aabb* (52-1-3) | 56.6 ± 4.31<sup>a</sup> | 3.94 ± 0.14<sup>b</sup> | 1.45 ± 0.07<sup>a</sup> | 44.7 ± 4.13<sup>a</sup> | 11.9 ± 0.71<sup>a</sup> |

Note: Values are means ± s.d. (n = 5 plants). Different letters at the same column indicate significant differences at $P < 0.05$ level.

### CRISPR/Cas9-induced mutations were transmitted to progeny

Only transgene-free T1 plants, derived from crossing the T0 mutants as maternal parents with a compatible WT plant, were examined to avoid complications that could have resulted from the continuing action of the CRISPR/Cas9 transgene. Among the 30 progeny of the primary mutant 52-1, 12 were transgene-free with ten of them carrying mutations. Meanwhile 3 out of 6 progeny of the cross between mutant 35-1 and a WT plant carried no transgenes and of these, only one carried mutations (Table 3). Additionally, 18 out of 20 progeny from the mutant 35-2 were transgene-free (Table 3), but only two carried mutation.

**Table 3.**
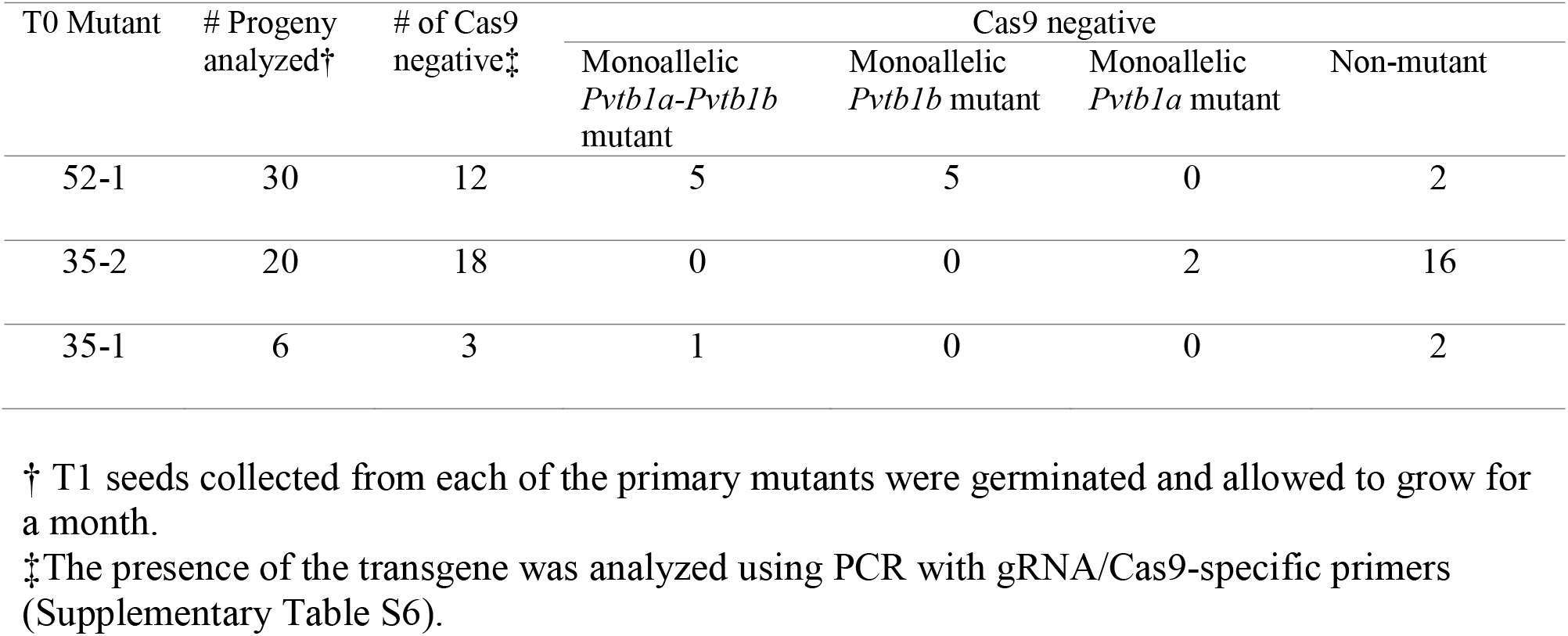
Transgene-free T1 mutants obtained from crossing each of the primary mutants with genetically compatible wild-type plants.

The *Pvtb1* mutations observed in all three primary mutants were also observed in their progeny, indicating a stable transmission of the original mutations to the progeny (Table S1). Seven T1 plants of the 52-1 had both *Pvtb1* genes mutated (Table S1), while the other five plants, in T1 generation, only contained mutations of *Pvtb1b* (Table S1). All mutations in these T1 plants were present in the primary mutant 52-1 except for that the 52-1-T1-24 has a mutated *Pvtb1a* allele carrying a 128 bp deletion which was not observed in the primary 52-1 mutant (Table S1). Two monoallelic *Pvtb1a* mutants (35-2-T1-1 and -6) were obtained from the progeny of the 35-2 (Table 3). Both carried identical mutations observed in the primary mutant 35-2 (Table S1). Furthermore, one doubly monoallelic *Pvtb1a-tb1b* mutant with big deletions for both *tb1* genes was found in the progeny of 35-1 (Table S1).

To test if mutant progeny still retains the phenotypic effects of *Pvtb1* mutations, tiller numbers of *Pvtb1b* monoallelic mutants (genotype AABb with A represents the WT and a represents the mutant allele of *Pvtb1a* and B represents the WT and b represents the mutant allele of *Pvtb1b*), and *Pvtb1a-Pvtb1b* doubly monoallelic mutant plants (genotype AaBb) in the T1 generation were investigated. Tiller numbers of doubly *Pvtb1a-Pvtb1b* monoallelic mutants and single *Pvtb1b* monoallelic mutants were significantly higher than the siblings carrying no mutations (null segregants), suggesting that *Pvtb1* genes act in dosage-dependent manner. The average percentage increase in tiller number was 45% for doubly *Pvtb1a-Pvtb1b* monoallelic mutant plants and 30% for the *Pvtb1b* monoallelic mutant plants (Table 4). Although the doubly monoallelic *Pvtb1a-Pvtb1b* mutant plants produced more tillers than the *Pvtb1b* monoallelic mutants, no significant difference was observed between the two groups of mutant plants, suggesting *Pvtb1b* plays a major role in tillering of switchgrass.

**Table 4.** The average tiller number of transgene-free T1 mutants and that of the siblings carrying no mutations (null segregants) and the percentage increase of tiller number for mutants over non-mutant siblings.

| Genotype | N | Average tiller number | % increase of tiller number in mutant over the WT |
| --- | --- | --- | --- |
| AABb (mutant) | 4 | $26.0 \pm 4.7^b$ | $30 \pm 23$ |
| AaBb (mutant) | 5 | $29.2 \pm 4.4^b$ | $45 \pm 20$ |
| AABB (WT) | 6 | $20.0 \pm 1.4^a$ | $0 \pm 10$ |

All data are shown as mean ± standard deviation. A and a represent the wild-type and mutant allele of *Pvtb1a* gene, respectively whereas B and b represent the wild-type and the mutant allele of the *Pvtb1b* gene, respectively. The percentage increase of tiller number for each genotype was calculated by comparing the mutant plants with the wild-type plants at the same developmental stage using the one-tailed Student's t-test. Different letters at the same column indicate significant differences at $P < 0.05$ level.

### mRNA-seq of *Pvtb1* mutants suggests *Pvtb1* genes are involved in multiple pathways that regulate tillering in switchgrass

Although the primary mutant 52-1 was later found to be chimeric, its enhanced tiller production (Liu et al., 2018) with 30% - 50% of the *tb1a* alleles and 61% - 94% of the *tb1b* alleles mutated are sufficient to be treated as a knockdown mutant of *Pvtb1* gene (Figure S3). Using a generalized linear model with the R package edgeR (Robinson et al., 2010), 831 genes showed significantly differential expression between the WT and 52-1 when the false discovery rate (FDR) was controlled at 0.05 (Benjamini and Hochberg, 1995) (Table S2). Among them, 364 genes were significantly upregulated while 467 genes showed significantly down regulation in the mutant 52-1 (Figure S4).

Orthologous Arabidopsis genes for these DEGs were subjected to Gene Ontology (GO) analysis using the Database for Annotation, Visualization and Integrated Discovery (DAVID) (Huang et al., 2009a; Huang et al., 2009b). Regarding the biological process, 27 terms (P<0.05, Table S4) were enriched for down-regulated genes in the mutant, and 12 terms (P<0.05, Table S3) for the up-regulated genes. A major GO term for these DEGs was DNA-templated transcription (GO: 0006351) (Figure 6A and 6B). Among the genes enriched in this GO term, transcript levels for 34 genes increased significantly in the *Pvtb1* gene knockdown mutant 52-1 while the remaining 43 genes were down-regulated (Table S5). The majority of the genes in this GO term encode transcription factors (TFs). For instance, genes for 3 TCP family TFs (TCP2, 4, and 5) and 4 MADS-box family TFs (AP1) were up-regulated in the mutant 52-1, while genes for 8 WRKY family TFs (WRKY30, 33, 40, 41, 46, and 53) and 3 NAC family TFs (NAC02, 036, and 102) were down-regulated. GO terms associated with cell differentiation and positive regulation of development were also enriched among the up-regulated genes in the mutant. Other GOs enriched in both up- and down-regulated genes were involved in response to various phytohormones suggesting *Pvtb1* genes are, as in other species involved in various hormonal signaling pathways in regulating tillering in switchgrass (Table S3 and S4). For example, BRH1, a brassinosteroid-responsive RING-H2 gene, was down-regulated in the *Pvtb1* gene knockdown mutant (Figure 7). In contrast, genes for auxin-responsive factor and ABA-induced proteins were both up-regulated in the mutant (Figure 7). In addition, GO terms involved in negative regulation of flower development and heterochromatin maintenance were overrepresented among the down-regulated genes (Figure 6B). For example, Arabidopsis DRD1 and EDM2 both regulate DNA methylation patterns (Cho et al., 2016; Duan et al., 2017) and the orthologs of these two were down-regulated in the present study. Hence, we speculate that epigenetic modification plays a role in *Pvtb1*-mediated regulation in tillering in switchgrass. Intriguingly, GOs associated with stress/defense responses were significantly enriched in down-regulated genes, indicating *Pvtb1* genes might function in maintaining homeostasis in the regulation of growth and defense (Figure 6B and Figure 7). These data suggest the gene regulatory network of tillering in switchgrass is complex and *Pvtb1* genes are important hubs in this network.

**Figure 6.**
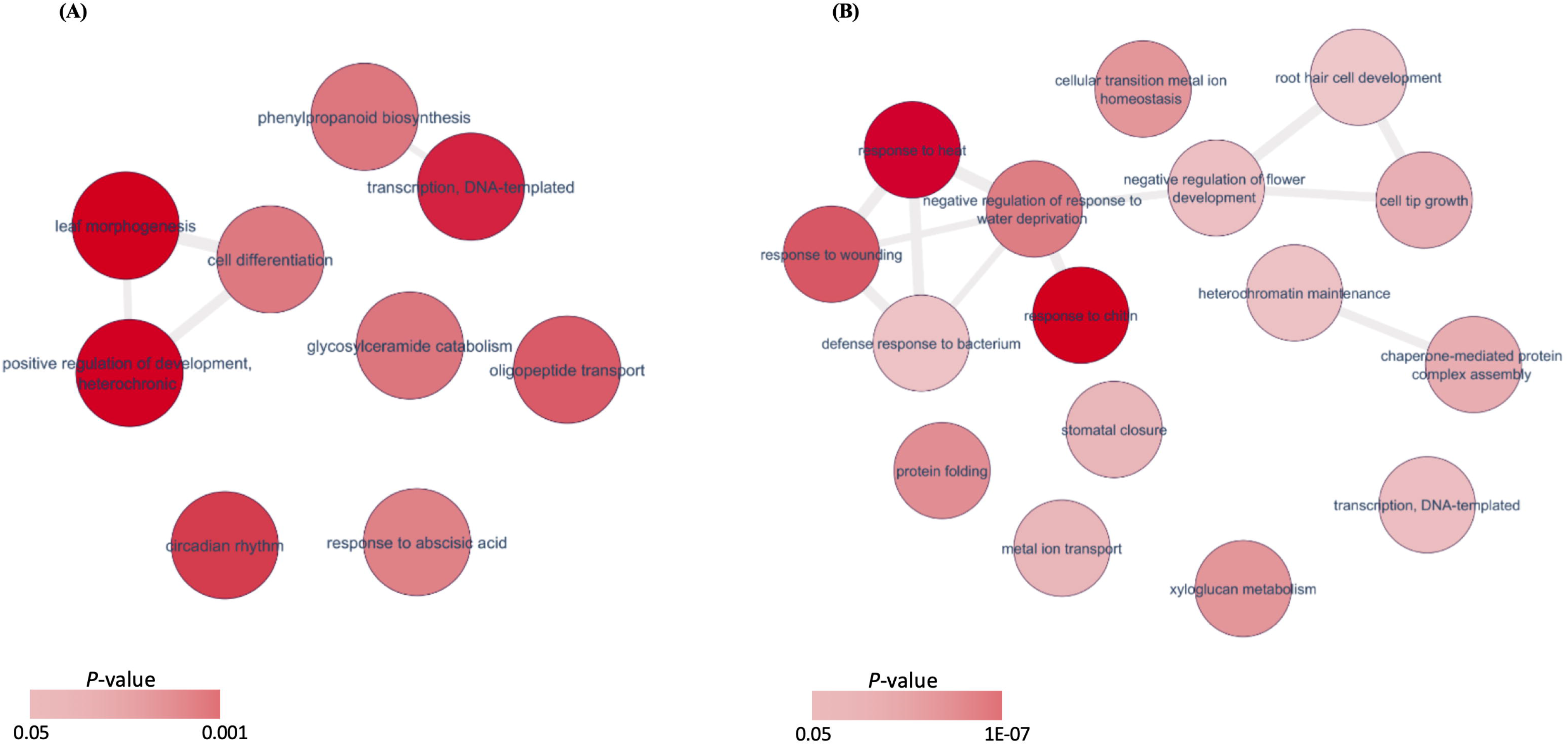
Enriched GO terms and differentially expressed genes of the *Pvtb1* genes knockdown mutant. A. Gene set enrichment analysis of up-regulated genes; B. Gene set enrichment analysis of down-regulated genes. Each red circle represents a GO term. Two color bars indicate *P-value* ranging from 0.05 to 0.001 and 1.0 × 10^−7^, respectively.

**Figure 7.**
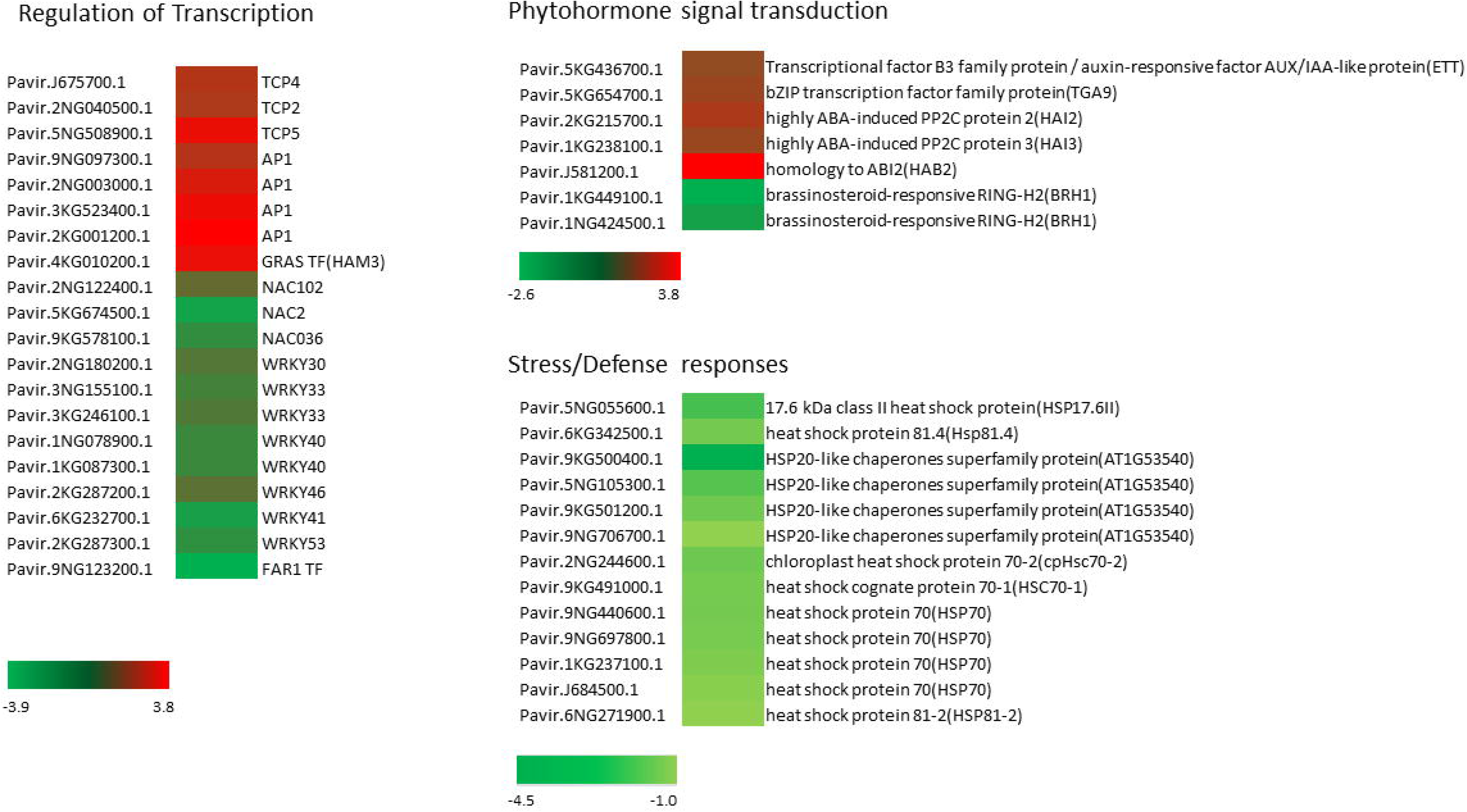
Expression patterns of representative differentially expressed genes (DEGs) involved in transcription regulation, phytohormone signal transduction and stress/defense responses previously characterized in Arabidopsis. Three color bars represent the range of fold change for DEGs, where red indicates up-regulation while green indicates down-regulation.

## Discussion

CRISPR/Cas9-based genome editing tool affords new possibilities for reverse genetics-based studies in switchgrass (Liu et al., 2018). In the present study, through the node culture of chimeric primary mutants induced by the CRISPR/Cas9 system, we successfully obtained a non-chimeric biallelic *Pvtb1a-Pvtb1b* mutant, and two non-chimeric mutants carrying biallelic *Pvtb1b* mutations and one *Pvtb1a* mutated allele. These results demonstrate that micropropagation is an effective way of isolating mutant sectors from chimeric mutants in switchgrass and obtaining non-chimeric mutants. For perennial grasses, it is difficult to obtain progeny seed due to the asynchronous flowering time and seed shattering (Cox et al., 2006). The low light intensity and lack of air movement in the greenhouse make it more difficult for container-grown transgenic switchgrass with insufficient root mass to produce viable seeds. Strong genetic-incompatibility of switchgrass is another obstacle to generate homozygous mutants. To set seeds carrying homozygous mutations, individual mutants used for crossing must have different alleles of S and Z genes (Martinez-Reyna and Vogel, 2002), which is difficult to determine because the molecular basis of the self-incompatibility in switchgrass has not been studied. Also, it takes over 6 months for switchgrass seedlings to reach reproductive stage (Hopkins et al., 1995; VanEsbroeck et al., 1997) for crossing. Given all these obstacles, the successful purification of chimeric mutants using micropropagation clearly has advantages.

Switchgrass cv. Alamo is an allotetraploid with two homeologous subgenomes, but detailed information about chromosome pairing, whole or partial genome duplications, and allelic diversity of specific genes is lacking (Missaoui et al., 2005; Okada et al., 2010). With limited information gained by sequencing clones of PCR amplicons spanning the target regions, the nature of mutants was not resolved unequivocally in our previous study (Liu et al., 2018). Using the Next Generation Sequencing (NGS) technology, genotypes of micropropagated mutants were fully characterized. The two primary mutants, 52-1 and 35-2 were both revealed by NGS to be chimeric mutants, which were likely the results of continuous action of CRISPR/Cas9 in somatic cells. Indeed, chimeric mutations induced by CRISPR/Cas9 have been reported in different species (Feng et al., 2014; Pan et al., 2017). For instance, Pan et al. (2017) showed that 63.9% T0 transgenic plants carried chimeric mutations in tomato (*Solanum lycopersicum* L.). In rice, the segregation ratio of CRISPR/Cas9-induced mutations in T1 generation did not follow the expected segregation ratio, indicating the chimeric mutations in T0 plants (Xu et al., 2015).

Transgene-free mutants were successfully obtained by crossing primary mutant plants with WT plants. The majority of the mutations observed in progeny were identical to the mutations in the primary mutants. For example, T1 progeny of the 52-1 carried the *Pvtb1b* mutations that were present in all five tillers of the primary mutant 52-1. These results demonstrate mutations induced by the CRISPR/Cas9 in T0 mutants were stably transmitted to the T1 generation without alteration. However, a mutation (128bp deletion) of *Pvtb1a*, not detected in the primary mutant 52-1, was observed in the T1 transgene-free mutant plant 52-1-T1-24. This deletion likely escaped detection in T0 plants due to the chimeric nature. The similar phenomenon has also been observed in maize (Lee et al., 2019).

*Pvtb1* genes are closely related to the *tb1* genes in other monocots. The *Pvtb1a-Pvtb1b* double biallelic mutants produced significantly higher tiller number than the WT plants under both hydroponic and soil culture conditions. Results from the hydroponic experiments in which starting plant materials were primary tillers validated the observed differences between the mutants and the WT plants were the result of genetic alteration instead of the existing physiological difference between the WT and the mutant. Our results strongly indicate *Pvtb1* genes play an important role in regulating tillering in switchgrass. There was no significant difference in the number of primary tillers between the WT and 52-1-3, suggesting the function of *Pvtb1* genes in switchgrass is only to regulate the rate of outgrowth of axillary buds that are destined to become tillers with one possible exception, i.e. the release of the outgrowth of the bud at the lowest node (Figure S2). This is similar to orthologs of *tb1* in other species, reflecting the functional conservation of *tb1* genes across species (Takeda et al., 2003; Kebrom et al., 2006; Aguilar-Martinez et al., 2007; Braun et al., 2012).

The *Pvtb1a-Pvtb1b* double biallelic mutant plants also produced more roots compared with WT plants. This result is consistent with Gaudin et al. (2014) who reports a decrease in *tb1* function in maize resulted in a larger root system. The increased root growth and development is likely the indirect result of increased tiller production, as each tiller normally develops its own adventitious roots. Increased tiller number has the potential to increase biomass yield in switchgrass (Chuck et al., 2011; Fu et al., 2012). The *Pvtb1a-Pvtb1b* mutants produced 29.6% and 15.5% more fresh and dry biomass, respectively. Although the difference on dry biomass between the wild-type and mutant plants is not statistically significant, this may change if mutant plants with more tillers and increased root mass are grown in the field where more resources are available.

In maize, studies have shown that *tb1* regulates branching in a dosage-dependent manner (Doebley et al., 1995; Hubbard et al., 2002). Maize heterozygous *tb1* mutants had slightly more tillers than WT plants, while homozygous *tb1* mutants produced more tillers than heterozygous *tb1* mutants (Doebley et al., 1995). In this study, we noticed the tiller number in monoallelic heterozygous *Pvtb1b* mutants (AABb) was significantly higher than WT plants, suggesting that *Pvtb1b* functions in a dosage-dependent manner. In addition, due to the high amino acid sequence identities between PvTB1A and PvTB1B, it is reasonable to expect they regulate tillering of switchgrass redundantly or additively. However, comparing monoallelic *Pvtb1b* mutant plants (AABb) with the doubly monoallelic *Pvtb1a-Pvtb1b* mutant plants (AaBb) did not show significant differences on tiller numbers, suggesting there is no significant additive effect between *Pvtb1a* and *Pvtb1b*. Therefore, *Pvtb1a* might have a minor effect on tillering in switchgrass. This is similar to Arabidopsis in which only BRC1 regulates branching, despite both BRC1 and BRC2 have the conserved TCP and R domains (Aguilar-Martinez et al., 2007; Gonzalez-Grandio et al., 2013; Seale et al., 2017).

To have a better understanding of the functions of *Pvtb1* genes, we examined global transcriptional changes caused by the down-regulation of *Pvtb1* genes. Increased expression level of genes for TCP TFs that are associated with cell differentiation and positive regulation of development in the mutant suggested that *Pvtb1* genes inhibit the tiller production through deactivating cell differentiation. In addition, *HAIRY MERISTEM 3* (*HAM3*), the gene for GRAS family TF that interacts with WUSCHEL (WUS) TF to promote shoot meristem development (Zhou et al., 2015), was up-regulated in the mutant, suggesting increased shoot stem cell proliferation in the *Pvtb1* gene knockdown mutant.

Our transcriptomic analysis results suggest PvTB1a and PvTB1b regulate tillering in switchgrass by interacting with complex hormonal signaling pathways. Six cytochrome P450 genes were up regulated in the mutant (Table S5). The members of cytochrome P450 family catalyze the biosynthesis of several phytohormones including auxin, brassinosteroids, and strigolactones which regulate branching across various plants species (Zhao, 2008; Kebrom et al., 2013). In addition, increased expression of ABA-responsive genes, *PROTEIN PHOSPHATASE 2C* (*PP2C*) genes, was observed in the mutant. These results suggest that *Pvtb1* genes regulate bud development by modulating phytohormone biosynthesis and signaling. In Arabidopsis and maize, it has been shown TB1/BRC1 promotes ABA accumulation and the expression of ABA response factors to inhibit bud outgrowth (Gonzalez-Grandio et al., 2013; Yao and Finlayson, 2015; Gonzalez-Grandio et al., 2017; Holalu and Finlayson, 2017; Dong, et al., 2019). Although it is well-known that *TB1*/*BRC1* are involved in hormonal signaling pathways in different plant species, these regulation pathways are not conserved across various species (Kebrom et al., 2013). For instance, cytokinins repress the expression of *TB1* in rice, while they act in a pathway independent of BRC1 in Arabidopsis (Aguilar-Martinez et al., 2007; Minakuchi et al., 2010). The timing of hormonal signaling in tillering has not been decided in switchgrass. Hence, more studies are needed to understand how the hormonal signals are involved in the *Pvtb1*-mediated regulation of branching.

Altered expression of genes in response to red or far-red light in the mutant suggest that *Pvtb1* genes integrate the light signal to regulate tillering in switchgrass (Figure 7). *FAR-RED-IMPAIRED RESPONSE1* (*FAR1*)-related sequence (FRS) family of transcription factors regulate plant growth and development in response to far-red light in Arabidopsis (Wang and Wang, 2015). The homolog of *FAR1*, *FAR-RED ELONGATED HYPOCOTYLS3* (*FHY3*) promotes shoot branching in Arabidopsis (Stirnberg et al., 2012). Further, *FAR1*/*FHY3* promotes *FHY1*/*FHL* gene expression to facilitate phyA nuclear accumulation under far-red light condition. The down-regulation of FRS TF genes in the switchgrass mutant suggested that *Pvtb1* genes may be involved in promotion of phyA nuclear accumulation to inhibit the axillary bud initiation or outgrowth. Additionally, knockdown of *Pvtb1* genes increased the expression level of the *Phytochrome interacting factor 4* (*PIF4*) gene that has been shown to regulate expression of genes involved in cell expansion (Huq and Quail, 2002). Because PIF4 is a TF regulated by phyB-mediated signaling, its activity is regulated by the red light signal (Xu, 2018). These results suggest *Pvtb1* genes regulate tillering through light signaling pathways. It has been reported that *tb1* genes inhibit bud outgrowth in the process of shade-avoidance-syndrome (SAS) (Kebrom et al., 2013). In Arabidopsis and sorghum, the expression levels of *BRC1* and *SbTB1* were both up-regulated under shade (Kebrom et al., 2006; Kebrom et al., 2010; Gonzalez-Grandio et al., 2013). Therefore, *Pvtb1* genes might also sense the low R:FR ratio to inhibit bud outgrowth in switchgrass.

Several genes associated with stress/defense responses were significantly down-regulated in the mutant. For example, 13 genes for Heat-shock proteins (Hsps)/chaperones which assist in protein refolding under stress conditions (Wang et al., 2004) were down-regulated in the mutant. It has been shown overexpression of alfalfa (*Medicago sativa* L.) *MsHSP70* gene could enhance Arabidopsis drought and heat stress tolerance (Li et al., 2017). Soybean (*Glycine max* (L.) Merril) Hsp 90 family members respond differentially to abiotic stresses and reduce the damage caused by abiotic stresses in Arabidopsis (Xu et al., 2013). Hence, the down-regulation of Hsp genes in the switchgrass mutant suggest hastened outgrowth of axillary buds of the mutant might trigger increased expression of stress/defense responsive genes. Further, the expression levels of several WRKY TF genes responsive to chitin elicitation functioning in plant defense to fungal pathogens (Libault et al., 2007) also decreased in the mutant. In Arabidopsis, WRKY TFs are necessary for resistance to pathogen infection (Zheng et al., 2006) or resistance to abiotic stresses (Chen et al., 2010). It is well-known tolerance-growth trade-offs occur in plants under the low-resource conditions (Koziol et al., 2012; Bristiel et al., 2018). It has been shown that TB1 may influences sucrose levels and energy balance within dormancy buds in maize (Dong, et al., 2019). Exploration of the mechanism of PvTB1s controlling energy balance in switchgrass would provide valuable information for the improvement of switchgrass for biomass production and development of enhanced stress-tolerant cultivars.

## Conclusions

We successfully isolated mutated segments from chimeric mutants using micropropagation. This method overcomes the difficulties of obtaining non-chimeric mutants in self-incompatible species. Further, transgene-free mutants were obtained in this research, which provided valuable germplasm for switchgrass genetic research and breeding. More importantly, we proved the stable transmission of mutations induced by the CRISPR/Cas9 system in switchgrass. We propose that *Pvtb1b* negatively regulates tillering in switchgrass, while the *Pvtb1a* may play a minor role on tillering. RNA-seq analysis revealed a complex regulatory network potentially regulating tillering in switchgrass and provided some clues to the pathways of *Pvtb1* genes.

## Acknowledgement

This work was partially supported by the National Institute of Food and Agriculture of the US Department of Agriculture (2013-33522-21091 to B.Y. and S.F.) and the Crop Bioengineering Center of Iowa State University (S.F.). The authors declare no competing financial interests and no conflict-of-interest. The authors also wish to thank Dr. Michael Baker at the ISU DNA Facility for his assistance on NGS experiment and Drs. Zhaoxia Li and Xianran Li for their valuable suggestions on NGS data analysis.

---

Supplementary Figure S1. Micropropagation of switchgrass. Longitudinally split nodal segments were cultured on the MS-0 medium without plant growth regulators.

Supplementary Figure S2. Outgrowth of the lowest axillary bud in mutant plants and the wild-type plants. Arrow indicates a tiller developing from the lowest node in the *Pvtb1a-Pvtb1b* mutant (52-1-3, aabb), which is usually absent in the WT. The WT plant was about 2 weeks older than the mutant plant. Other tillers were removed for better view.

Supplementary Figure S3. Estimation of allelic composition of *Pvtb1* genes in cDNA samples of the mutant 52-1.

Supplementary Figure S4. Mean-Difference plot showing the log-fold change and average abundance of each gene. Significantly up- and down-regulated genes in the mutant are highlighted in red and blue, respectively.

## References

Aguilar-Martinez J, Poza-Carrion C, Cubas P (2007) Arabidopsis BRANCHED1 acts as an integrator of branching signals within axillary buds. Plant Cell 19: 458–472

Alexandrova K, Denchev P, Conger B (1996) Micropropagation of switchgrass by node culture. Crop Science 36: 1709–1711

Anders S, Pyl P, Huber W (2015) HTSeq-a Python framework to work with high-throughput sequencing data. Bioinformatics 31: 166–169

Arite T, Iwata H, Ohshima K, Maekawa M, Nakajima M, Kojima M, Sakakibara H, Kyozuka J (2007) DWARF10, an RMS1/MAX4/DAD1 ortholog, controls lateral bud outgrowth in rice. Plant Journal 51: 1019–1029

Benjamini Y, Hochberg Y (1995) Controlling the False Discovery Rate: A Practical and Powerful Approach to Multiple Testing. Journal of the Royal Statistical Society Series B-Methodological 57: 289–300

Boe A (2007) Variation between two switchgrass cultivars for components of vegetative and seed biomass. Crop Science 47: 636–642

Boe A, Beck D (2008) Yield components of biomass in switchgrass. Crop Science 48: 1306–1311

Bolger A, Lohse M, Usadel B (2014) Trimmomatic: a flexible trimmer for Illumina sequence data. Bioinformatics 30: 2114–2120

Braun N, de Saint Germain A, Pillot J, Boutet-Mercey S, Dalmais M, Antoniadi I, Li X, Maia-Grondard A, Le Signor C, Bouteiller N, Luo D, Bendahmane A, Turnbull C, Rameau C (2012) The Pea TCP Transcription Factor PsBRC1 Acts Downstream of Strigolactones to Control Shoot Branching. Plant Physiology 158: 225–238

Bristiel P, Gillespie L, Ostrem L, Balachowski J, Violle C, Volaire F (2018) Experimental evaluation of the robustness of the growth-stress tolerance trade-off within the perennial grass Dactylis glomerata. Functional Ecology 32: 1944–1958

Casler M (2010) Changes in Mean and Genetic Variance During Two Cycles of Within-family Selection in Switchgrass. Bioenergy Research 3: 47–54

Chen H, Lai Z, Shi J, Xiao Y, Chen Z, Xu X (2010) Roles of arabidopsis WRKY18, WRKY40 and WRKY60 transcription factors in plant responses to abscisic acid and abiotic stress. Bmc Plant Biology 10

Cho EJ, Choi SH, Kim JH, Kim JE, Lee MH, Chung BY, Woo HR, Kim J-H (2016) A mutation in plant-specific SWI2/SNF2-like chromatin-remodeling proteins, DRD1 and DDM1, delays leaf senescence in Arabidopsis thaliana. PloS one 11: e0146826

Choi M, Woo M, Koh E, Lee J, Ham T, Seo H, Koh H (2012) Teosinte Branched 1 modulates tillering in rice plants. Plant Cell Reports 31: 57–65

Chuck G, Tobias C, Sun L, Kraemer F, Li C, Dibble D, Arora R, Bragg J, Vogel J, Singh S, Simmons B, Pauly M, Hake S (2011) Overexpression of the maize Corngrass1 microRNA prevents flowering, improves digestibility, and increases starch content of switchgrass. Proceedings of the National Academy of Sciences of the United States of America 108: 17550–17555.

Cox T, Glover J, Van Tassel D, Cox C, DeHaan L (2006) Prospects for developing perennial-grain crops. Bioscience 56: 649–659

Das A, Gosal S, Sidhu J, Dhaliwal H (2000) Induction of mutations for heat tolerance in potato by using in vitro culture and radiation. Euphytica 114: 205–209

Das M, Fuentes R, Taliaferro C (2003) Genetic variability and trait relationships in switchgrass. Crop Science 44: 443–448

Davidson D, Chevalier P (1987) Influence of Polyethylene Glycol-Induced Water Deficits on Tiller Production in Spring Wheat 1. Crop science 27: 1185–1187

Dixon L, Greenwood J, Bencivenga S, Zhang P, Cockram J, Mellers G, Ramm K, Cavanagh C, Swain S, Boden S (2018) TEOSINTE BRANCHED1 Regulates Inflorescence Architecture and Development in Bread Wheat (Triticum aestivum). Plant Cell 30: 563–581

Dobin A, Davis C, Schlesinger F, Drenkow J, Zaleski C, Jha S, Batut P, Chaisson M, Gingeras T (2013) STAR: ultrafast universal RNA-seq aligner. Bioinformatics 29: 15–21

Doebley J, Stec A, Gustus C (1995) teosinte branched1 and the origin of maize: evidence for epistasis and the evolution of dominance. Genetics 141: 333–346

Doebley J, Stec A, Hubbard L (1997) The evolution of apical dominance in maize. Nature 386: 485–488

Dong, Z., Xiao, Y., Govindarajulu, R., Feil, R., Siddoway, M.L., Nielsen, T., Lunn, J.E., Hawkins, J., Whipple, C. and Chuck, G (2019) The regulatory landscape of a core maize domestication module controlling bud dormancy and growth repression. Nature communications, 10(1): 1–15.

Duan C-G, Wang X, Zhang L, Xiong X, Zhang Z, Tang K, Pan L, Hsu C-C, Xu H, Tao WA (2017) A protein complex regulates RNA processing of intronic heterochromatin-containing genes in Arabidopsis. Proceedings of the National Academy of Sciences 114: E7377–E7384

Feng Z, Mao Y, Xu N, Zhang B, Wei P, Yang D, Wang Z, Zhang Z, Zheng R, Yang L, Zeng L, Liu X, Zhu J (2014) Multigeneration analysis reveals the inheritance, specificity, and patterns of CRISPR/Cas-induced gene modifications in Arabidopsis. Proceedings of the National Academy of Sciences of the United States of America 111: 4632–4637

Finlayson S, Krishnareddy S, Kebrom T, Casal J (2010) Phytochrome Regulation of Branching in Arabidopsis. Plant Physiology 152: 1914–1927

Fu C, Sunkar R, Zhou C, Shen H, Zhang J, Matts J, Wolf J, Mann D, Stewart C, Tang Y, Wang Z (2012) Overexpression of miR156 in switchgrass (Panicum virgatum L.) results in various morphological alterations and leads to improved biomass production. Plant Biotechnology Journal 10: 443–452

Gaudin AC, McClymont SA, Soliman SS, Raizada MN (2014) The effect of altered dosage of a mutant allele of Teosinte branched 1 (tb1-ref) on the root system of modern maize. BMC genetics 15: 23

Gonzalez-Grandio E, Pajoro A, Franco-Zorrilla J, Tarancon C, Immink R, Cubas P (2017) Abscisic acid signaling is controlled by a BRANCHED1/HD-ZIP I cascade in Arabidopsis axillary buds. Proceedings of the National Academy of Sciences of the United States of America 114: E245–E254

Gonzalez-Grandio E, Poza-Carrion C, Sorzano C, Cubas P (2013) BRANCHED1 Promotes Axillary Bud Dormancy in Response to Shade in Arabidopsis. Plant Cell 25: 834–850

Goodstein D, Shu S, Howson R, Neupane R, Hayes R, Fazo J, Mitros T, Dirks W, Hellsten U, Putnam N, Rokhsar D (2012) Phytozome: a comparative platform for green plant genomics. Nucleic Acids Research 40: D1178–D1186

Harten AM, Bouter H, Broertjes C (1981) In vitro adventitious bud techniques for vegetative propagation and mutation breeding of potato (*Solanum tuberosum* L.). II. Significance for mutation breeding. Euphytica 30: 1–8

He Y, Zhu M, Wang L, Wu J, Wang Q, Wang R, Zhao Y (2018) Programmed self-elimination of the CRISPR/Cas9 construct greatly accelerates the isolation of edited and transgene-free rice plants. Molecular plant 11: 1210–1213

Holalu S, Finlayson S (2017) The ratio of red light to far red light alters Arabidopsis axillary bud growth and abscisic acid signalling before stem auxin changes. Journal of Experimental Botany 68: 943–952

Hopkins AA, Vogel KP, Moore KJ, Johnson KD, Carlson IT (1995) Genotype Effects and Genotype by Environment Interactions for Traits of Elite Switchgrass Populations. Crop Science 35: 125–132

Huang D, Sherman B, Lempicki R (2009a) Bioinformatics enrichment tools: paths toward the comprehensive functional analysis of large gene lists. Nucleic Acids Research 37: 1–13

Huang D, Sherman B, Lempicki R (2009b) Systematic and integrative analysis of large gene lists using DAVID bioinformatics resources. Nature Protocols 4: 44–57

Hubbard L, McSteen P, Doebley J, Hake S (2002) Expression patterns and mutant phenotype of teosinte branched1 correlate with growth suppression in maize and teosinte. Genetics 162: 1927–1935

Huq E, Quail P (2002) PIF4, a phytochrome-interacting bHLH factor, functions as a negative regulator of phytochrome B signaling in Arabidopsis. Embo Journal 21: 2441–2450

Jiang W, Yang B, Weeks D (2014) Efficient CRISPR/Cas9-Mediated Gene Editing in Arabidopsis thaliana and Inheritance of Modified Genes in the T2 and T3 Generations. Plos One 9

Kebrom T, Brutnell T, Finlayson S (2010) Suppression of sorghum axillary bud outgrowth by shade, phyB and defoliation signalling pathways. Plant Cell and Environment 33: 48–58

Kebrom T, Burson B, Finlayson S (2006) Phytochrome B represses Teosinte Branched1 expression and induces sorghum axillary bud outgrowth in response to light signals. Plant Physiology 140: 1109–1117

Kebrom T, Spielmeyer W, Finnegan E (2013) Grasses provide new insights into regulation of shoot branching. Trends in Plant Science 18: 41–48

Koziol L, Rieseberg LH, Kane N, Bever JD (2012) Reduced drought tolerance during domestication and the evolution of weediness results from tolerance–growth trade‐offs. Evolution: International Journal of Organic Evolution 66: 3803–3814

Lee K, Zhang Y., Kleinstiver B.P., Guo J.A., Aryee M.J., Miller J., Malzahn A., Zarecor S., Lawrence-Dill, C.J., Joung, J.K., Qi, Y., Wang K (2019) Activities and specificities of CRISPR/Cas9 and Cas12a nucleases for targeted mutagenesis in maize. Plant biotechnology journal 17: 362–372.

Li Z, Long R, Zhang T, Wang Z, Zhang F, Yang Q, Kang J, Sun Y (2017) Molecular cloning and functional analysis of the drought tolerance gene MsHSP70 from alfalfa (Medicago sativa L.). Journal of Plant Research 130: 387–396

Libault M, Wan J, Czechowski T, Udvardi M, Stacey G (2007) Identification of 118 Arabidopsis transcription factor and 30 ubiquitin-ligase genes responding to chitin, a plant-defense elicitor. Molecular Plant-Microbe Interactions 20: 900–911

Liu Y, Merrick P, Zhang Z, Ji C, Yang B, Fei Sz (2018) Targeted mutagenesis in tetraploid switchgrass (Panicum virgatum L.) using CRISPR/Cas9. Plant biotechnology journal 16: 381–393

Lu F, Lipka A, Glaubitz J, Elshire R, Cherney J, Casler M, Buckler E, Costich D (2013) Switchgrass Genomic Diversity, Ploidy, and Evolution: Novel Insights from a Network-Based SNP Discovery Protocol. Plos Genetics 9

Maluszynski M, Ahloowalia BS, Sigurbjörnsson B (1995) Application of *in vivo* and *in vitro* mutation techniques for crop improvement. Euphytica 85: 303–315

Martin-Trillo M, Cubas P (2010) TCP genes: a family snapshot ten years later. Trends in Plant Science 15: 31–39

Martin-Trillo M, Grandio E, Serra F, Marcel F, Rodriguez-Buey M, Schmitz G, Theres K, Bendahmane A, Dopazo H, Cubas P (2011) Role of tomato BRANCHED1-like genes in the control of shoot branching. Plant Journal 67: 701–714

Martinez-Reyna J, Vogel K (2002) Incompatibility systems in switchgrass. Crop Science 42: 1800–1805

McLaughlin S, Kszos L (2005) Development of switchgrass (Panicum virgatum) as a bioenergy feedstock in the United States. Biomass & Bioenergy 28: 515–535

Minakuchi K, Kameoka H, Yasuno N, Umehara M, Luo L, Kobayashi K, Hanada A, Ueno K, Asami T, Yamaguchi S, Kyozuka J (2010) FINE CULM1 (FC1) Works Downstream of Strigolactones to Inhibit the Outgrowth of Axillary Buds in Rice. Plant and Cell Physiology 51: 1127–1135

Missaoui A, Paterson A, Bouton J (2005) Investigation of genomic organization in switchgrass (Panicum virgatum L.) using DNA markers. Theoretical and Applied Genetics 110: 1372–1383.

Mitchell R, Schmer M (2012) Switchgrass harvest and storage. In Switchgrass. Springer, pp 113–127

Mitchell R, Vogel K, Sarath G (2008) Managing and enhancing switchgrass as a bioenergy feedstock. Biofuels Bioproducts & Biorefining-Biofpr 2: 530–539

Morris D (1977) Transport of exogenous auxin in two-branched dwarf pea seedlings (*Pisum sativum* L.). Planta 136: 91–96

Narasimhamoorthy B, Saha M, Swaller T, Bouton J (2008) Genetic Diversity in Switchgrass Collections Assessed by EST-SSR Markers. Bioenergy Research 1: 136–146

Nicolas M, Rodriguez-Buey M, Franco-Zorrilla J, Cubas P (2015) A Recently Evolved Alternative Splice Site in the BRANCHED1a Gene Controls Potato Plant Architecture. Current Biology 25: 1799–1809

Novák FJ, Afza R, Vanduren M, Omar M (1990) Mutation induction by gamma irradiation of in *vitro cult*ured shoot-tips of banana and plantain (Musa *cvs)*. Tropical Agriculture 67: 21–28

Okada M, Lanzatella C, Saha M, Bouton J, Wu R, Tobias C (2010) Complete Switchgrass Genetic Maps Reveal Subgenome Collinearity, Preferential Pairing and Multilocus Interactions. Genetics 185: 745–760

Pan C, Ye L, Qin L, Liu X, He Y, Wang J, Chen L, Lu G (2017) CRISPR/Cas9-mediated efficient and heritable targeted mutagenesis in tomato plants in the first and later generations (vol 6, 24765, 2016). Scientific Reports 7

Pyott DE, Sheehan E, Molnar A (2016) Engineering of CRISPR/Cas9‐mediated potyvirus resistance in transgene‐free Arabidopsis plants. Molecular plant pathology 17: 1276–1288

Rameau C, Bertheloot J, Leduc N, Andrieu B, Foucher F, Sakr S (2015) Multiple pathways regulate shoot branching. Frontiers in Plant Science 5

Reddy S, Finlayson S (2014) Phytochrome B Promotes Branching in Arabidopsis by Suppressing Auxin Signaling. Plant Physiology 164: 1542–1550

Robinson M, McCarthy D, Smyth G (2010) edgeR: a Bioconductor package for differential expression analysis of digital gene expression data. Bioinformatics 26: 139–140

Seale M, Bennett T, Leyser O (2017) BRC1 expression regulates bud activation potential but is not necessary or sufficient for bud growth inhibition in Arabidopsis. Development 144: 1661–1673

Shannon P, Markiel A, Ozier O, Baliga N, Wang J, Ramage D, Amin N, Schwikowski B, Ideker T (2003) Cytoscape: A software environment for integrated models of biomolecular interaction networks. Genome Research 13: 2498–2504

Sievers F, Wilm A, Dineen D, Gibson T, Karplus K, Li W, Lopez R, McWilliam H, Remmert M, Soding J, Thompson J, Higgins D (2011) Fast, scalable generation of high-quality protein multiple sequence alignments using Clustal Omega. Molecular Systems Biology 7

Sorefan K, Booker J, Haurogne K, Goussot M, Bainbridge K, Foo E, Chatfield S, Ward S, Beveridge C, Rameau C, Leyser O (2003) MAX4 and RMS1 are orthologous dioxygenase-like genes that regulate shoot branching in Arabidopsis and pea. Genes & Development 17: 1469–1474

Stirnberg P, Zhao S, Williamson L, Ward S, Leyser O (2012) FHY3 promotes shoot branching and stress tolerance in Arabidopsis in an AXR1-dependent manner. Plant Journal 71: 907–920

Studer A, Zhao Q, Ross-Ibarra J, Doebley J (2011) Identification of a functional transposon insertion in the maize domestication gene tb1. Nature genetics 43: 1160

Supek F, Bosnjak M, Skunca N, Smuc T (2011) REVIGO Summarizes and Visualizes Long Lists of Gene Ontology Terms. Plos One 6

Takeda T, Suwa Y, Suzuki M, Kitano H, Ueguchi-Tanaka M, Ashikari M, Matsuoka M, Ueguchi C (2003) The OsTB1 gene negatively regulates lateral branching in rice. Plant Journal 33: 513–520

Thimann K, Skoog F (1933) Studies on the growth hormone of plants III The inhibiting action of the growth substance on bud development. Proceedings of the National Academy of Sciences of the United States of America 19: 714–716

VanEsbroeck G, Hussey M, Sanderson M (1997) Leaf appearance rate and final leaf number of switchgrass cultivars. Crop Science 37: 864–870

Waltz E (2018) With a free pass, CRISPR-edited plants reach market in record time. Nature Biotechnology 36: 6–7

Wang H, Wang H (2015) Multifaceted roles of FHY3 and FAR1 in light signaling and beyond. Trends in Plant Science 20: 453–461

Wang M, Le Moigne M-A, Bertheloot J, Crespel L, Perez-Garcia M-D, Ogé L, Demotes-Mainard S, Hamama L, Davière J-M, Sakr S (2019) BRANCHED1: a key hub of shoot branching. Frontiers in plant science 10

Wang W, Simmonds J, Pan Q, Davidson D, He F, Battal A, Akhunova A, Trick HN, Uauy C, Akhunov E (2018) Gene editing and mutagenesis reveal inter-cultivar differences and additivity in the contribution of *TaGW2* homoeologues to grain size and weight in wheat. In. Springer Berlin Heidelberg, Theoretical and Applied Genetics, pp 1–13

Wang W, Vinocur B, Shoseyov O, Altman A (2004) Role of plant heat-shock proteins and molecular chaperones in the abiotic stress response. Trends in Plant Science 9: 244–252

Wang X, Tilford C, Neuhaus I, Mintier G, Guo Q, Feder J, Kirov S (2017) CRISPR-DAV: CRISPR NGS data analysis and visualization pipeline. Bioinformatics 33: 3811–3812

Whipple C, Kebrom T, Weber A, Yang F, Hall D, Meeley R, Schmidt R, Doebley J, Brutnell T, Jackson D (2011) grassy tillers1 promotes apical dominance in maize and responds to shade signals in the grasses. Proceedings of the National Academy of Sciences of the United States of America 108: E506–E512

Wright L, Turhollow A (2010) Switchgrass selection as a “model” bioenergy crop: A history of the process. Biomass & Bioenergy 34: 851–868.

Xu D (2018) Multifaceted Roles of PIF4 in Plants. Trends in Plant Science 23: 749–751

Xu J, Xue C, Xue D, Zhao J, Gai J, Guo N, Xing H (2013) Overexpression of GmHsp90s, a Heat Shock Protein 90 (Hsp90) Gene Family Cloning from Soybean, Decrease Damage of Abiotic Stresses in Arabidopsis thaliana. Plos One 8

Xu R, Li H, Qin R, Li J, Qiu C, Yang Y, Ma H, Li L, Wei P, Yang J (2015) Generation of inheritable and “transgene clean” targeted genome-modified rice in later generations using the CRISPR/Cas9 system. Scientific Reports 5

Yang J, Luo D, Yang B, Frommer W, Eom J (2018) SWEET11 and 15 as key players in seed filling in rice. New Phytologist 218: 604–615

Yao C, Finlayson S (2015) Abscisic Acid Is a General Negative Regulator of Arabidopsis Axillary Bud Growth. Plant Physiology 169: 611–626

Zhang Y, Liang Z, Zong Y, Wang Y, Liu J, Chen K, Qiu J-L, Gao C (2016) Efficient and transgene-free genome editing in wheat through transient expression of CRISPR/Cas9 DNA or RNA. Nature communications 7: 12617

Zhang Y, Zalapa J, Jakubowski A, Price D, Acharya A, Wei Y, Brummer E, Kaeppler S, Casler M (2011) Natural Hybrids and Gene Flow between Upland and Lowland Switchgrass. Crop Science 51: 2626–2641

Zhao C, Zhang Z, Xie S, Si T, Li Y, Zhu J (2016) Mutational Evidence for the Critical Role of CBF Transcription Factors in Cold Acclimation in Arabidopsis. Plant Physiology 171: 2744–2759

Zhao Y (2008) The role of local biosynthesis of auxin and cytokinin in plant development. Current Opinion in Plant Biology 11: 16–22

Zheng Z, Abu Qamar S, Chen Z, Mengiste T (2006) Arabidopsis WRKY33 transcription factor is required for resistance to necrotrophic fungal pathogens. Plant Journal 48: 592–605

Zhou Y, Liu X, Engstrom E, Nimchuk Z, Pruneda-Paz J, Tarr P, Yan A, Kay S, Meyerowitz E (2015) Control of plant stem cell function by conserved interacting transcriptional regulators. Nature 517: 377–U528

Zou J, Zhang S, Zhang W, Li G, Chen Z, Zhai W, Zhao X, Pan X, Xie Q, Zhu L (2006) The rice HIGH-TILLERING DWARF1 encoding an ortholog of Arabidopsis MAX3 is required for negative regulation of the outgrowth of axillary buds. Plant Journal 48: 687–696

